# Nuclear m6A Reader YTHDC1 Promotes Muscle Stem Cell Activation/Proliferation by Regulating mRNA Splicing and Nuclear Export

**DOI:** 10.1101/2022.08.07.503064

**Authors:** Yulong Qiao, Qiang Sun, Xiaona Chen, Di Wang, Ruibao Su, Yuanchao Xue, Hao Sun, Huating Wang

**Affiliations:** Department of Orthopaedics and Traumatology, Li Ka Shing Institute of Health Sciences, The Chinese University of Hong Kong, Hong Kong, China; Department of Chemical Pathology, Li Ka Shing Institute of Health Sciences, The Chinese University of Hong Kong, Hong Kong, China; Key Laboratory of RNA Biology, Institute of Biophysics, Chinese Academy of Sciences, Beijing, China; Center for Neuromusculoskeletal Restorative Medicine (CNRM), CUHK InnoHK Centres, The Chinese University of Hong Kong

**Keywords:** Muscle stem cell, Muscle regeneration, YTHDC1, m6A, hnRNPG

## Abstract

Skeletal muscle stem cells (also known as satellite cells, SCs) are essential for muscle regeneration and the regenerative activities of SCs are intrinsically governed by gene regulatory mechanisms but the post-transcriptional regulation in SCs remains largely unknown. N(6)-methyladenosine (m6A) modification of RNAs is the most pervasive and highly conserved RNA modification in eukaryotic cells and exerts powerful impact on almost all aspects of mRNA processing which is mainly endowed by its binding with m6A reader proteins. Here in this study, we investigate the previously uncharacterized regulatory roles of YTHDC1, a m6A reader in SCs. Our results demonstrate YTHDC1 is an essential regulator of SC activation and proliferation upon acute injury induced muscle regeneration. The induction of YTHDC1 is indispensable for SC activation and proliferation thus inducible YTHDC1 depletion almost abolishes SC regenerative capacity. Mechanistically, transcriptome-wide profiling using LACE-seq in both SCs and C2C12 myoblasts identifies m6A mediated binding targets of YTHDC1. Next, splicing analysis defines splicing mRNA targets of m6A-YTHDC1. Furthermore, nuclear export analysis also leads to identification of potential mRNA export targets of m6A-YTHDC1 in SCs and C2C12 myoblasts and interestingly some mRNAs can be regulated at both splicing and export levels. Lastly, we map YTHDC1 interacting protein partners in myoblasts and unveil a myriad of factors governing mRNA splicing, nuclear export and transcription, among which hnRNPG appears to be a bona fide interacting partner of YTHDC1. Altogether, our findings uncover YTHDC1 as an essential factor controlling SC regenerative ability through multi-faceted gene regulatory mechanisms in myoblast cells.

## Introduction

Skeletal muscle has a robust regenerative capacity, with rapid re-establishment of full power occurring even after severe damage that causes widespread myofiber necrosis. This is accomplished by SCs which normally lie quiescent and uniquely labeled by the expression of paired box (Pax) transcription factor Pax7^1^. Upon injury, the master myogenic regulator MyoD is expressed to enable rapid activation of SCs, which then expand as proliferating myoblasts; the myoblasts then differentiate and fuse into myofibers to repair the damaged muscle, while a subset self-renews to restore the quiescent SC pool. Deregulated SC activity contributes to the development of many muscle diseases; it is thus imperative to understand the way SCs contribute to regeneration. Intrinsically, both transcriptional and post-transcriptional gene regulations constitute key mechanisms governing SC activities^2, 3^. Here in this study we investigate the post-transcriptional gene regulation mediated by N(6)-methyladenosine (m6A) modification.

m6A is the most pervasive and highly conserved RNA modification in eukaryotic cells, and exerts powerful impact on almost all aspects of mRNA processing including alternative splicing, nuclear export, stability maintenance and translational efficiency, thus regulating diverse cellular processes^4^. The past decade has witnessed rapid technical and conceptual advances that allowed transcriptome-wide interrogation of m6A dynamics and galvanized interest in m6A function. It is widely accepted^4, 5^ that RNA m6A modification is catalyzed by a multicomponent methyltransferase complex (the “writer”) comprising METTL14, METTL3 and WTAP etc., and can be removed by specific demethylases (the “erasers”) including FTO and ALKBH5. However, it is the so called “reader” proteins that bind m6A and impart the epitranscriptomic information engraved in RNA m6A to functional signals, thus these proteins are considered as key effectors and executors of m6A function. So far, a category of m6A readers has been identified and classified as several families, among which YTH domain family proteins, YTHDF1, YTHDF2, YTHDF3, YTHDC1 and YTHDC2, specifically recognize m6A sites through YTH domain and are the most well-known readers.

YTHDC1 is a unique m6A reader because of its dominant location in the nucleus endowing its post-transcriptional regulatory functions such as pre-mRNA splicing^6, 7^, mRNA export^8, 9^ and mRNA stabilization^10–12^. The pioneer study from Xiao W et. al. ^6^ show that YTHDC1 promotes exon inclusion of targeted mRNAs in 293T cells through recruiting pre-mRNA splicing factor SRSF3 while blocking SRSF10 binding^6^; however, in a separate study^9^, YTHDC1 interacting with SRSF3 facilitates the delivery of methylated mRNAs to export receptor NXF1 for cytoplasmic export, pointing to YTHDC1 and SRSF3 as adaptor proteins coupling mRNA splicing and export^9^. Interestingly, rapidly evolving evidence from the past two years implicates YTHDC1 in epigenetic control^13, 14^. This function arises from its ability to bind chromatin associated RNAs (caRNAs), followed by recruitment of various histone modifying factors such as SETDB1^15^, KAP1^16^, or KDM3B^17^ to gate chromatin accessibility and downstream transcription. Beyond epigenetic control, recent studies also show that YTHDC1 participates in transcriptional regulation through promoting RNA polymerase II pausing release^18^, facilitating transcriptional condensates formation and gene activation of eRNAs^19^; moreover, it can stimulate nascent RNA synthesis by inhibiting integrator termination complex^20^. There is thus a current advent in illuminating molecular underpinnings of m6A dependent functions of YTHDC1. Key questions remain: is splicing regulation the predominant mechanism underlying YTHDC1 function in any cell? Can YTHDC1 exert its function via multiple mechanisms in any given cell type? Are there undiscovered transcriptional or post-transcriptional regulatory mechanisms of m6A-YTHDC1? In this project, we elucidate m6A-YTHDC1 functional mechanisms in skeletal muscle stem cells and muscle regeneration. Surprisingly, until now the study of m6A regulation and function in SCs is scarce ^21–23^ and it remains to be answered: what are the functional readers in SCs? Does YTHDC1 control SC activities and how?

Our findings in this study identify YTHDC1 as an essential regulator of SC activation/proliferation thus the acute injury induced muscle regeneration. Inducible YTHDC1 knockout impairs SC activation and proliferation, which blocks muscle regeneration. Mechanistically, combining transcriptome-wide YTHDC1 binding profiles and global m6A map we define m6A-YTHDC1 regulatory targets in myoblasts. Further analyses demonstrate YTHDC1 loss alters both splicing and nuclear export of target mRNAs. Lastly, we uncover that YTHDC1 interacts with a myriad of proteins in proliferating myoblasts including regulators of mRNA splicing, mRNA nuclear export as well as transcriptional regulators, strengthening its diverse gene regulatory functions. Altogether, our findings suggest that YTHDC1 is a previously uncharacterized factor governing SC activities and muscle regeneration and it can exert pleiotropic gene regulatory functions through interacting with different proteins in myoblast cells.

## Results

### m6A regulators are dynamically expressed during SC lineage progression and YTHDC1 is induced upon SC activation/proliferation

To investigate possible roles of m6A regulators in SCs, we analyzed the expression dynamics of a panel of m6A writers, readers and erasers using our recently generated RNA-seq datasets^24^ from SCs at various time points of lineage progression (Fig. 1A). Freshly isolated SCs (FISCs) by fluorescence-activated cell sorting (FACS) sorting, are considered as early activating cells due to the isolation process^25^; quiescent SCs (QSCs) were obtained through a pre-fixation step with paraformaldehyde (PFA) before the FACS sorting to preserve the quiescence status; FISCs were cultured for 24, 48 and 72 hours (h) to obtain fully activated, proliferating and differentiating SCs (ASC-24h, −48h, and −72h). As a result, we found that many m6A regulators were dynamically expressed in the lineage progression course (Fig. 1B-C and Suppl. Fig. S1), reflecting their possible importance in controlling different phases of SC activities. Specifically, we found the reader *Ythdc1* mRNA was expressed at all stages with the highest level in FISCs (Fig. 1B-D). By Western blotting (WB) and immunofluorescence (IF) (Fig. 1E, F), we observed a very low level of YTHDC1 protein in FISC and an evident increase of YTHDC1 protein in activating cells (ASC-24h) and continued increase in proliferating cells (ASC-48h); YTHDF1 and YTHDF2 reader proteins appeared to show similar expression dynamics. Consistent with prior studies^6, 26^, YTHDC1 protein was predominantly detected in the nucleus but not cytoplasm (Fig. 1F). This was also confirmed in mouse C2C12 cell line (a commonly used surrogate for ASCs) by fractionation assay (Fig. 1G), suggesting its possible nuclear regulatory functions in myoblast cells.

**Fig. 1.**
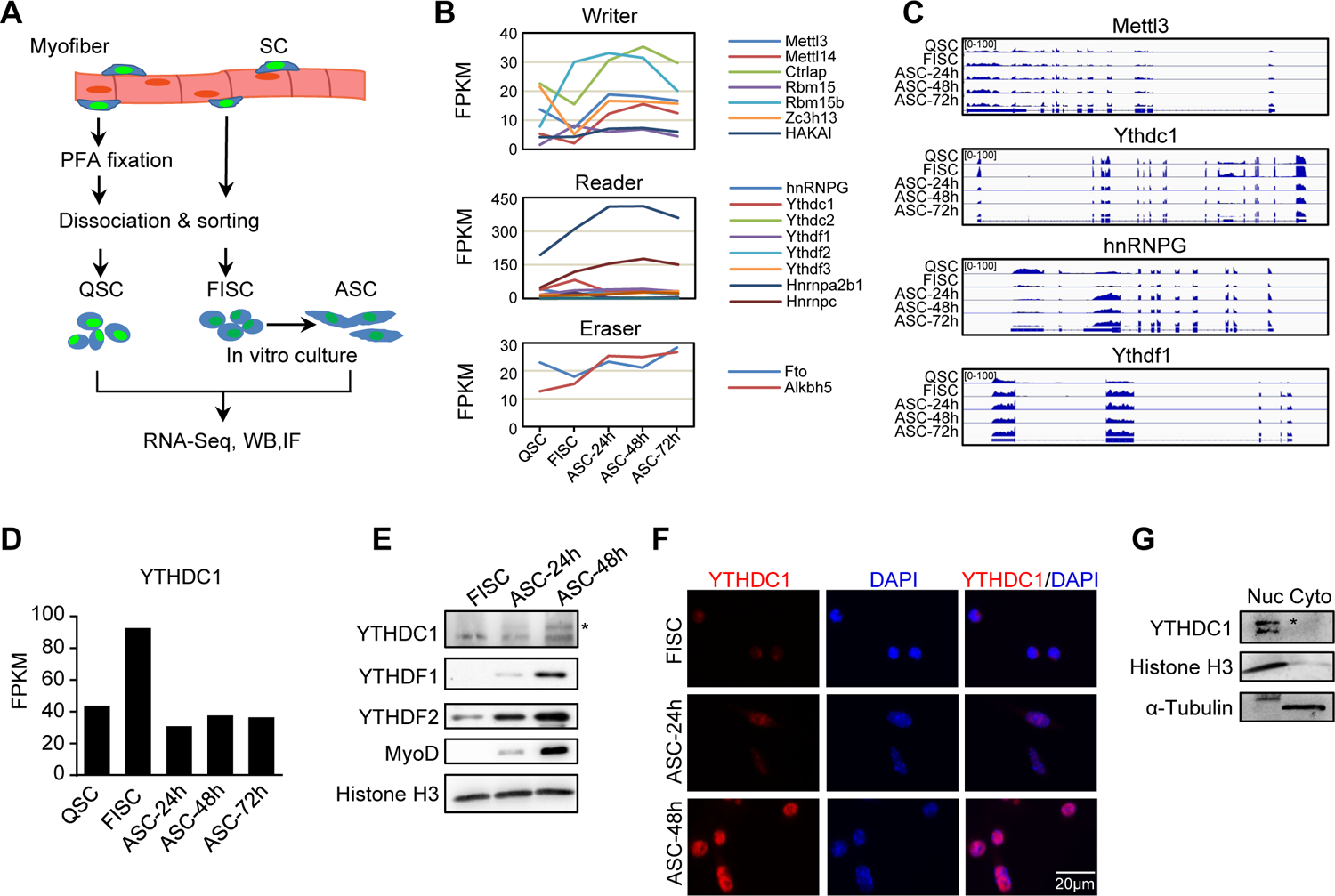
m6A regulators are dynamically expressed during SC lineage progression and YTHDC1 is induced upon SC activation/proliferation. A. Schematic illustration of SC collection from Pax7-nGFP mice. Fixed quiescent SCs (QSC), freshly isolated SCs (FISC) and cultured SCs (ASC) were subject to RNA-seq, western blotting (WB), and Immunofluorescence (IF) analyses. B. The expression dynamics of m6A writer, reader and eraser proteins in the above cells from analyzing the RNA-seq data. C. Representative RNA-seq tracks showing the expression dynamics of the selected m6A regulators. D. The expression dynamic of YTHDC1 mRNA (FPKM) from RNA-seq. E. WB showing the induction of YTHDC1, YTHDF1 and YTHDF2 proteins upon SC activation and proliferation. *denotes the correct position of YTHDC1. Histone H3 was used as a loading control. F. IF staining showing the induction of YTHDC1 protein upon SC activation and proliferation. Scale bar=20μm. G. WB showing the predominant location of YTHDC1 in nuclear portion of C2C12 myoblasts.

### Inducible YTHDC1 deletion in SCs abolishes acute injury induced muscle regeneration

Despite intensive investigation of YTHDC1 regulatory mechanisms, genetic evidence remains largely lacking to support its roles in biological processes. To test its biological function in SCs, we crossed a recently available *Ythdc1* floxed allele^7^ with the *Pax7^CreER^*;*ROSA^EYFP^* mouse^27^ to generate an inducible knock out (iKO) mouse to inactivate YTHDC1 specifically in SCs (Fig. 2A). After successful removal of YTHDC1 by five consecutive plus two extra doses of Tamoxifen (TMX) (Fig. 2B-C), BaCl_2_ was injected into the tibialis anterior (TA) muscle to induce acute damage^3^. In control (Ctrl) littermate mice, the regeneration followed a predictable course^3^. Massive immune cell infiltration was observed on the first day post injury (dpi) (data not shown); meanwhile, SCs were rapidly activated and reached a peak of proliferation at 3dpi (data not shown); by 5dpi they were mainly differentiating and labeled with eMyHC+ (expressed only in newly regenerating fibers) (Fig. 2D-E), coinciding with the initiation of myofiber repair. At 14dpi, injured muscles were largely repaired (Suppl. Fig. 2A-C). As a striking contrast, muscle regeneration was nearly completely abolished in the iKO muscles; excessive immune infiltration was still present at 5 or 7 dpi with no signs of repair (Fig. 2D); a sharp decrease of eMyHC+ myofibers was detected at both time points (Fig. 2E); and a complete loss of Pax7+ cells (Fig. 2F-G) was also observed despite no difference on uninjured muscles (Fig. 2F-G). The regeneration was in fact never observed even at 28dpi; iKO muscle remained significantly smaller than Ctrl (Suppl. Fig. S2A-C). Altogether, the above data indicate the essential function of YTHDC1 in acute injury induced muscle regeneration.

**Fig. 2.**
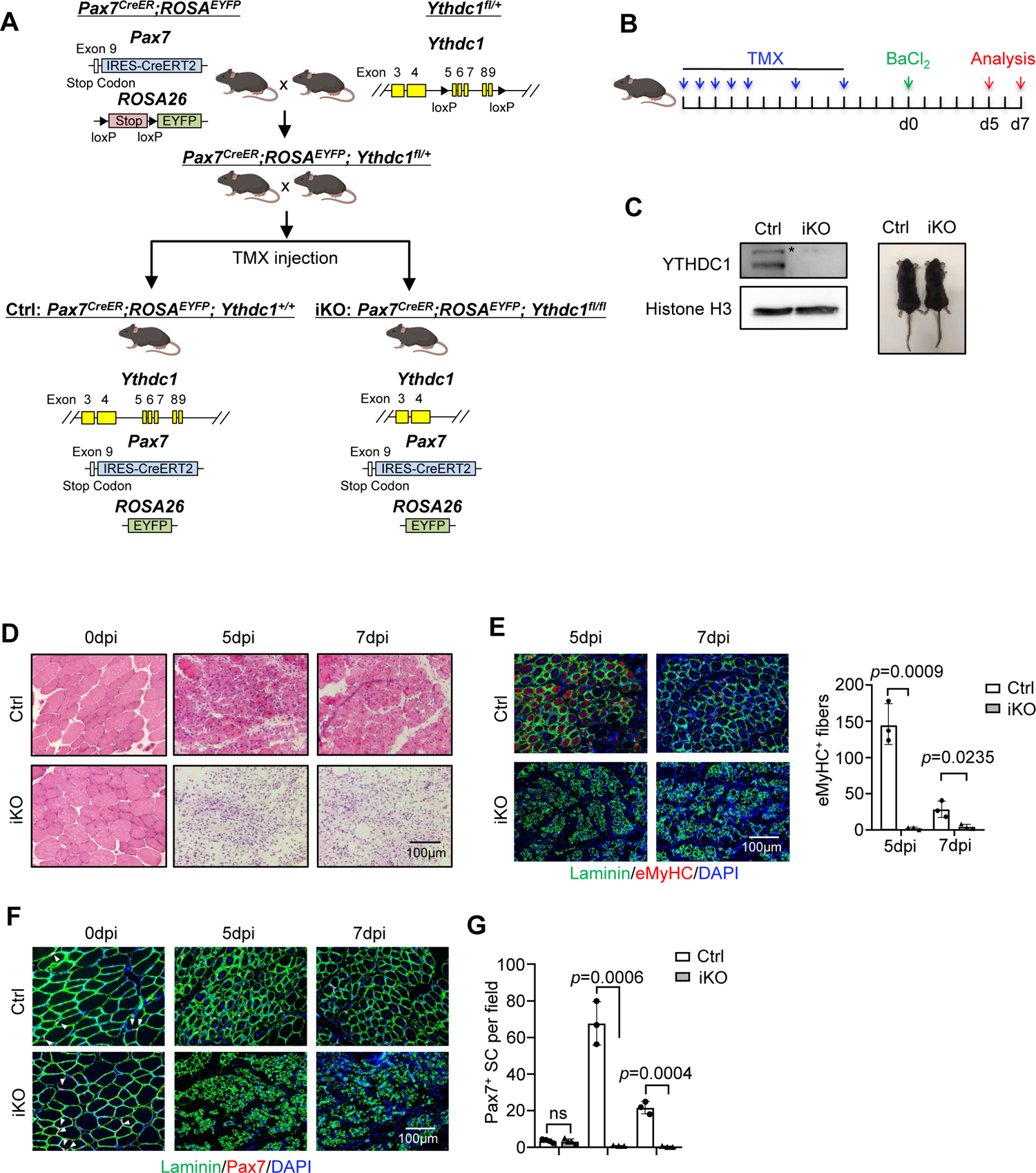
Inducible YTHDC1 deletion in SCs abolishes acute injury induced muscle regeneration. **A.** Breeding scheme for generating YTHDC1 inducible knockout (iKO) and control (Ctrl) mice. **B.** Schematic outline of the tamoxifen (TMX) administration used in the study and experimental design for testing the effect of YTHDC1 deletion on barium chloride (BaCl_2_) induced muscle regeneration process. **C.** Left: WB showing the deletion of YTHDC1 in ASC-48h from iKO but not Ctrl mice. Right: no obvious morphological difference was detected in iKO vs. Ctrl mice. **D.** H&E staining of the above injured muscles at 0, 5 and 7 dpi. Scale bar=100μm. **E.** Left: Immunostaining of eMyHC (red) and laminin (green) of the above injured TA muscles at 5 and 7 dpi. Scale bar=100μm. Right: Quantification of eMyHC positive fibers per field. *n*=3 mice per group. **F.** Immunostaining of Pax7 (red) and laminin (green) on TA muscle sections at 0, 5 and 7 dpi. Scale bar=100μm. **G.** Quantification of Pax7 positive SCs per field at 0, 5 and 7 dpi. *n*=4 mice per group for 0dpi, *n*=3 mice per group for 5 and 7dpi. Bars represent mean ± s.d. for all graphs. Statistical significance was determined using a two-tailed Student’s t test.

### Inducible YTHDC1 knockout impairs SC activation/proliferation

To pinpoint the major defects of iKO SCs that block the regeneration, we suspected YTHDC1 loss impaired SC activation and proliferation considering it was induced upon SC activation and highly expressed in proliferating myoblasts (Fig. 1D-E). To this end, FISCs from Ctrl or iKO mice were cultured for 24 or 48h; and indeed the iKO cells displayed evident growth arrest (Suppl. Fig. S3A-B). This was confirmed by 4h EdU treatment and staining, activation (24h) and proliferation (48h) was drastically inhibited in the iKO compared to the Ctrl (Fig. 3A). On isolated single myofibers, SCs also failed to proliferate by EdU staining of the fibers cultured for 48h (Fig. 3B). Consistently, staining for Pax7 and MyoD showed that a significantly lower percentage of double positive cells were detected at both ASC-24h and 48h in the iKO compared to the Ctrl (Fig. 3C); the protein levels of Pax7 and MyoD were also largely diminished in the iKO cells at 48h (Suppl. Fig. S3C). On single myofibers, the number of Pax7+ MyoD+ SCs were also significantly reduced at both 24h and 48h (Fig. 3D). Lastly, *in vivo* EdU assay was also performed (Fig. 3E). EdU was injected 2 days after BaCl_2_ induced injury and SCs were isolated 12h later for staining. A significant reduction of EdU+ cells was observed in the iKO compared to the Ctrl muscles (Fig. 3F). Altogether, the above findings demonstrate YTHDC1 loss causes a severe defect in SC activation and proliferation thus solidify the essential function of YTHDC1 in early stages of SC regenerative activities.

**Fig. 3.**
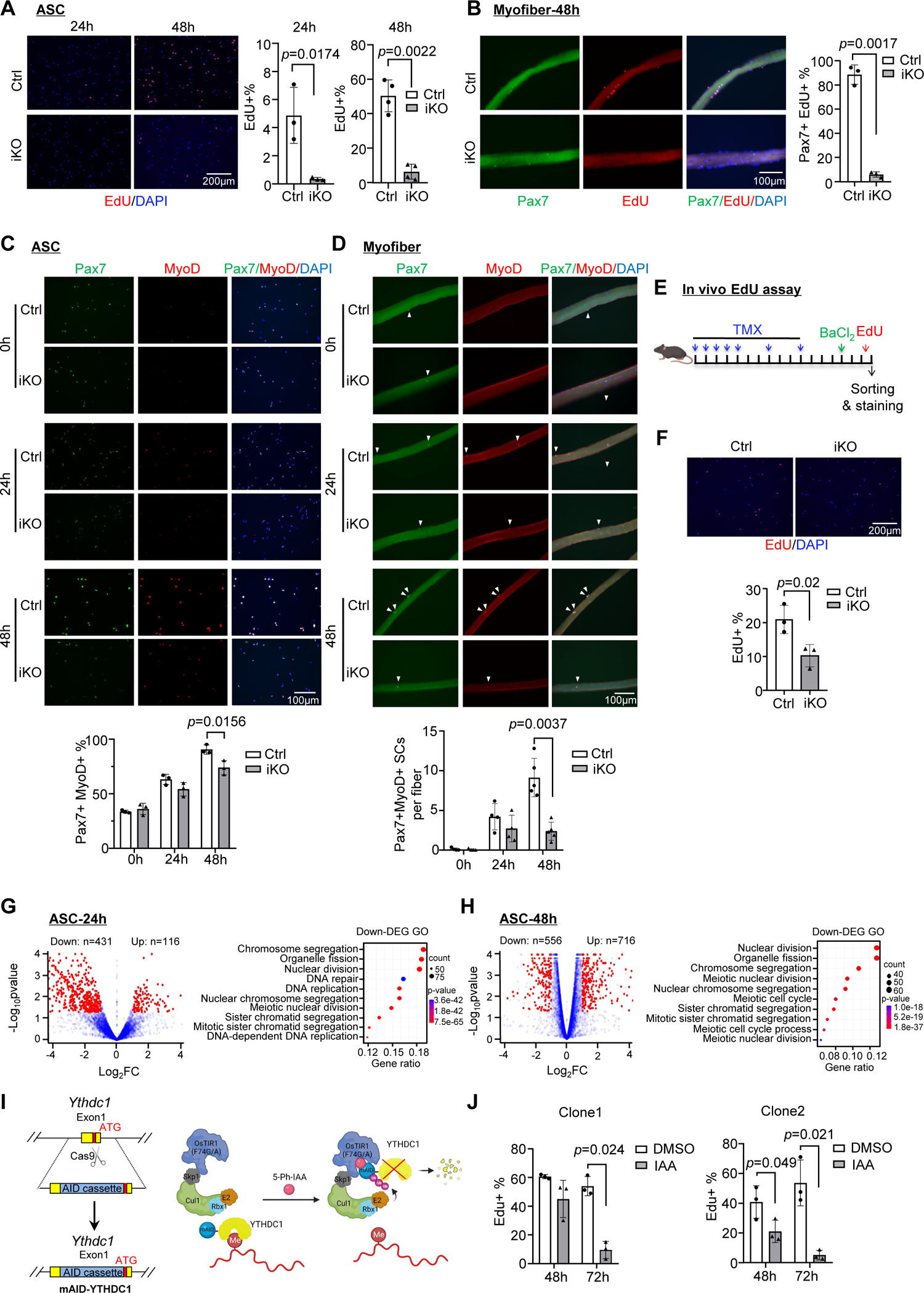
Inducible YTHDC1 knockout impairs SC activation/proliferation. **A.** Left: EdU (red) staining of ASC-24h and ASC-48h from iKO and Ctrl mice. Scale bar=200μm. Right: Quantification of percentage of EdU+ cells. *n*=3 mice per group for ASC-24h, *n*=4 mice per group for ASC-48h. **B.** Left: EdU (red) staining of EDL myofibers isolated from Ctrl or iKO and cultured for 48h. Scale bar=100μm. Right: Quantification of the percentage of Pax7+EdU+ SCs. *n*=3 mice per group. **C.** Top: IF staining of Pax7 (green) and MyoD (red) of FISC (0h), ASC-24h and ASC-48h from Ctrl and iKO mice. Scale bar=100μm. Bottom: Quantification of the percentage of Pax7+MyoD+ cells. *n*=3 mice per group. **D.** Top: IF staining of Pax7 (green) and MyoD (red) on EDL myofibers at 0h (freshly isolated), 24h and 48h. Scale bar=100μm. Bottom: Quantification of the number of Pax7+MyoD+ cells per fiber. (*n*=4 mice per group for 0 and 24h, *n*=5 mice per group for 48h) **E.** Schematic illustration of *in vivo* EdU assay in Ctrl and iKO mice. **F.** Top: EdU staining of the above freshly isolated and fixed SCs at 3 dpi. Scale bar=200μm. Bottom: Quantification of the percentage of EdU+ cells in iKO vs. Ctrl. *n*=3 mice per group. **G-H**. Left: RNA-seq was performed in ASC-24h or −48h from iKO and Ctrl. Volcano plot showing the down- and up-regulated genes in iKO vs. Ctrl. Right: GO analysis for the down-regulated genes. **I.** Schematic illustration of generating a C2C12 cell line with inducible YTHDC1 degradation using the auxin-inducible degron (AID2) system. **J.** Two independent mAID-YTHDC1 cell lines were treated with DMSO or 5-Ph-IAA(IAA) for the indicated time. EdU assay was performed and the percentage of EdU+ cells were quantified at the designated time points. *n*=3 replicates. Bars represent mean ± s.d. for all graphs. Statistical significance was determined using a two-tailed Student’s t test.

Next, to gain more insight, transcriptomic analyses using RNA-seq was conducted in ASC-24h cells. As a result, a total of 547 differentially expression genes (DEGs) were identified in iKO vs. Ctrl SCs with 431 down- and 116 up-regulated (Fig. 3G and Suppl. Table S1); the down-regulated genes were enriched for GO terms such as “Nuclear division”, “Chromosome segregation” etc. (Fig. 3G and Suppl. Table S1). When the assay was performed in ASC-48h cells, a total of 1272 DEGs (556 down- and 716 up-regulated) were identified (Fig. 3H) and again the down-regulated genes were enriched for similar GO terms as the above (Fig. 3H and Suppl. Table S1). These findings are in accordance with the activation/proliferation defect observed in the iKO cells.

Lastly, we generated a C2C12 mouse myoblast cell line with inducible YTHDC1 degradation using the auxin-inducible degron (AID2) system^28^ (Fig. 3I). An mAID-mCherry tag was knocked into the *Ythdc1* N-terminal locus in a C2C12 cell line expressing auxin receptor Osir1 (F74G) (Fig. 3I). As expected, the addition of 5-Ph-IAA (auxin, IAA) but not DMSO (negative control) induced rapid degradation of the mAID tagged YTHDC1 (Suppl. Fig. S3D-E). In line with the results obtained from ASCs, decreased proliferation rate was observed in two independent AID-YTHDC1 clones treated with IAA for both 48h or 72h (Suppl. Fig. S3F and Fig. 3J), strengthening that YTDHC1 functions to promote myoblast proliferation.

### LACE-seq defines transcriptome-wide YTHDC1 binding profiles in myoblasts

To fathom the underlying mechanism, we conducted transcriptome-wide binding analysis for YTHDC1, reasoning ultimately it is the binding sites/targets that determine its function. To this end, we harnessed the recently developed LACE-seq (the Linear Amplification of Complementary DNA Ends and Sequencing) method (Fig. 4A) which enables global profiling of RNA-binding protein (RBP) target sites with a relatively low quantity of starting cellular material^29^. Around one million of ASC-48h were subject to LACE-seq with two biological replicates (B1, B2) and four technical replicates (T1, T2) (Fig. 4B). As a result, a total of 2444 shared peaks corresponding to 951 genes were identified from comparing the 4 replicates and defined as YTHDC1 targets (Fig. 4B and Suppl. Table S2). All of these replicates were well correlated with each other (Pearson correlation coefficient: 0.82∼0.91). These genes were enriched for GO terms such as “Cell projection organization” and “Cell junction assembly” etc. (Fig. 4C and Suppl. Table S2). To reinforce the result, we also performed LACE-seq with C2C12 myoblasts; two technical replicates were included and a much higher number of cells (10 million) were used for each replicate since there was no difficulty growing and obtaining C2C12 cells (Fig. 4A-B).

**Fig. 4.**
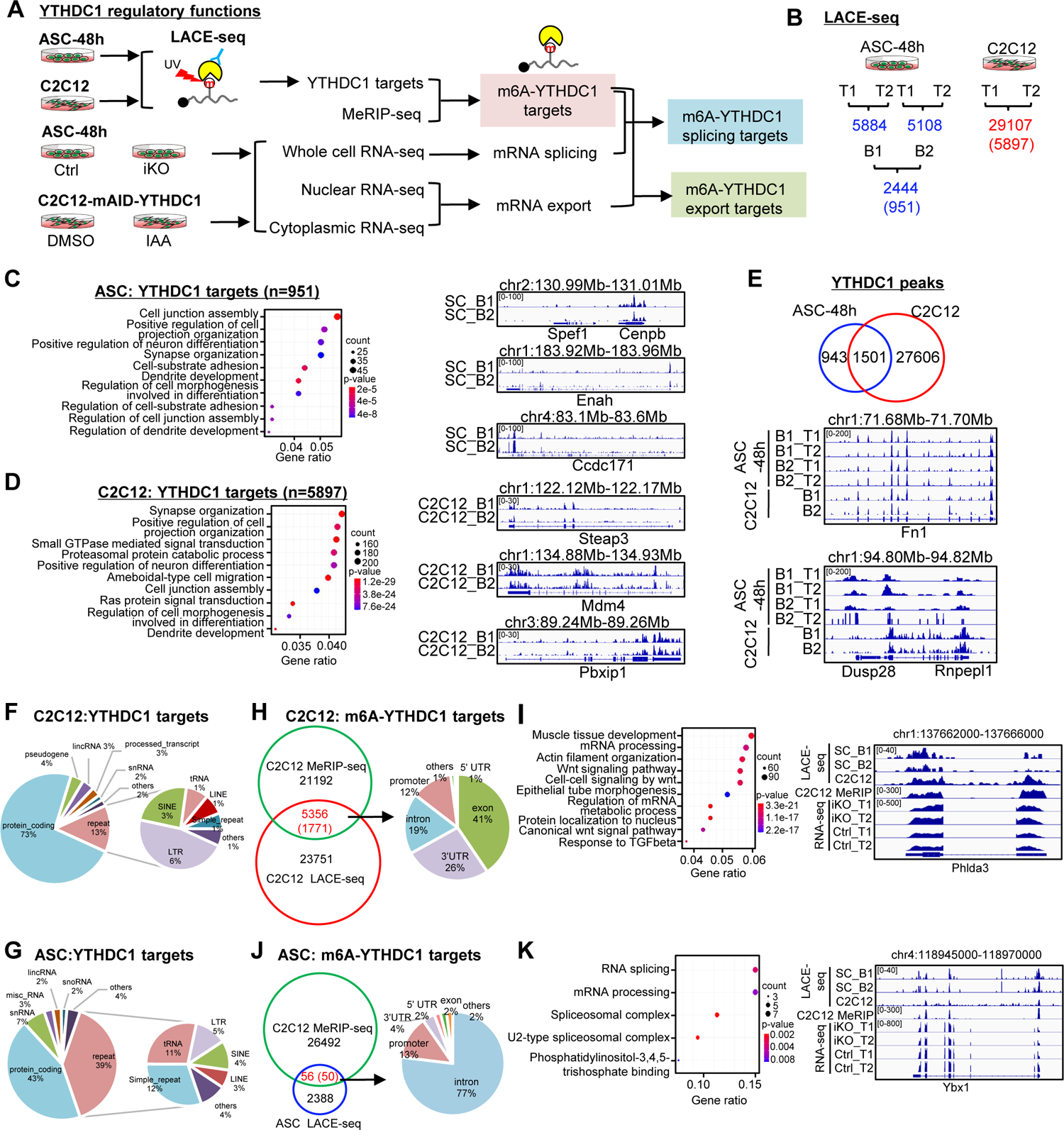
LACE-seq defines transcriptome-wide YTHDC1 binding profiles in myoblasts. **A.** Schematic illustration of the experimental design for performing LACE-seq and subsequent combination with MeRIP-seq, bulk RNA-seq, subcellular RNA-seq for defining and elucidating YTHDC1 splicing/export targets and post-transcriptional regulation. **B.** LACE-seq was performed in both ASC-48h and C2C12 myoblasts and the number of identified peaks in each technical (T) or biological replicate (B) and the shared number of peaks (genes) between the replicates are shown. **C.** Left: Go analysis for the identified YTHDC1 targets in ASCs. Right: genome tracks for three selected genes. **D.** Left: Go analysis for the identified YTHDC1 targets in C2C12. Right: genome tracks for three selected genes. **E.** Top: Overlapping between the above identified C2C12 and ASC peaks. Bottom: Genome tracks for two selected genes. **F-G.** Left: the genome distribution of YTHDC1 binding peaks in C2C12 or ASC. Right: detailed distribution of the YTHDC1 binding peaks on repeat regions. **H.** Left: Integrating C2C12 MeRIP-seq data with the C2C12 LACE-seq identified 5356 regions (1771 mRNAs) as m6A-YTHDC1 targets. Right: Distribution of YTHDC1 binding on the above targets. **I.** Left: GO analysis of the above 1771 targets. Right: Genomic tracks of a selected target, Phlda3. **J.** Left: Integrating the C2C12 MeRIP-seq data with the ASC LACE-seq identified 56 regions (50 mRNAs) as m6A-YTHDC1 targets. Right: Distribution of YTHDC1 binding on the above identified target mRNAs. **K.** Left: GO analysis of the above 50 targets. Right: Genomic tracks of a selected target, Ybx1.

Our results showed that data from C2C12 was of very high quality with the two replicates displaying high correlation (Pearson correlation coefficient: 0.96). A total of 29107 shared peaks corresponding to 5897 genes were identified as YTHDC1 binding targets (Fig. 4B and Suppl. Table S2). Very similar to the above result from ASCs, these genes were also enriched for a variety of GO terms such as “Cell projection organization” and “Cell junction assembly” etc. (Fig. 4D and Suppl. Table S2). In fact, 1501 out of the 2444 identified peaks in ASCs could be found in C2C12 (Fig. 4E and Suppl. Table S2), testifying the success of the assay. Next, further probing into the binding locations, we found that in both C2C12 (Fig. 4F and Suppl. Table S2) and ASCs (Fig. 4G and Suppl. Table S2), a large portion of the peaks (73% and 43%) were mapped to protein coding genes and small portions in lincRNAs, snRNAs, snoRNAs etc. Interestingly, a significant portion was mapped to repeat regions such as SINE (13%) and LINE (39%), which was consistent with prior reports^15^. Overall, these findings are largely consistent with prior reports from conducting YTHDC1 global profiling through CLIP-seq or RIP-seq^6, 15, 16, 30^, suggesting regulating mRNA processing is probably the dominant role of YTHDC1 in mouse myoblasts.

To further tease out m6A dependent function of YTHDC1, we obtained the published methylated RNA immunoprecipitation sequencing (MeRIP-seq) data from C2C12^31^ which was of decent quality judged by the enrichment of a classical RRACH motif in the identified binding sites and also dominant enrichment at 3’UTR genomic regions (data not shown). By integrating the dataset with the above C2C12 LACE-seq dataset, a total of 5356 peaks (1771 genes) were identified; 41% of the peaks resided in exons followed by 3’UTRS (26%), introns (19%) and promoters (12%) (Fig. 4H and Suppl. Table S2). The above defined final set of 1771 m6A-YTHDC1 targets were enriched for variable GO terms (Fig. 4I and Suppl. Table S2), among which Wnt signaling and TGFβ pathways are related to myoblast proliferation (Fig. 4I and Suppl. Table S2). Next, we also compared the ASC LACE-seq data with the above C2C12 MeRIP-seq data and identified a total of 56 peaks (corresponding to 50 genes) (Fig. 4J and Suppl. Table S2); interestingly, these genes were highly enriched for GO terms such as “RNA splicing”, “mRNA processing” etc. (Fig. 4K and Suppl. Table S2) which were also found in C2C12 (Fig. 4I), suggesting m6A-YTHDC1 not only modulates RNA processing but also controls the processing of RNA processing factors.

### YTHDC1 depletion in ASCs leads to altered splicing events

Next we tested if YTHDC1 plays a role in modulating mRNA splicing as splicing regulation is its best known function^6, 7^. To this end, we analyzed the splicing events and altered events upon YTHDC1 depletion using the RNA-seq data from Fig. 3G-H (Fig. 5A). Expectedly, in both ASC-24h and −48h, a high number of differential splicing events (DSEs) were uncovered in the iKO compared to the Ctrl cells (Fig. 5A); skipped exons (SEs) and mutually exclusive exons (MXEs) dominated these events, which was consistent with previously reported role for YHTDC1 in promoting exon inclusion^6^. Among the 951 targets of YTHDC1, a total of 189 displayed DSE (Fig. 5B and Suppl. Table S3), thus defined as YTHDC1 splicing targets. These targets were enriched for “Regulation of actin filament-based process”, “Dendrite development” (Suppl. Fig. S4A and Suppl. Table S3), among which we experimentally validated the altered splicing on Palb2, Scn5a, Lrp8 mRNAs by RT-PCR assay (Fig. 5C). To pinpoint m6A-YTHDC1 dependent splicing events, we then intercepted the DSEs with the 50 m6A-YTHDC1 targets identified in ASCs (Fig. 4J) and 11 of them were defined as m6A-YTHDC1 splicing targets (Fig. 5D), among which the altered splicing on Itbp3bp and Nek1 mRNAs was experimentally validated (Fig. 5E).

**Fig. 5.**
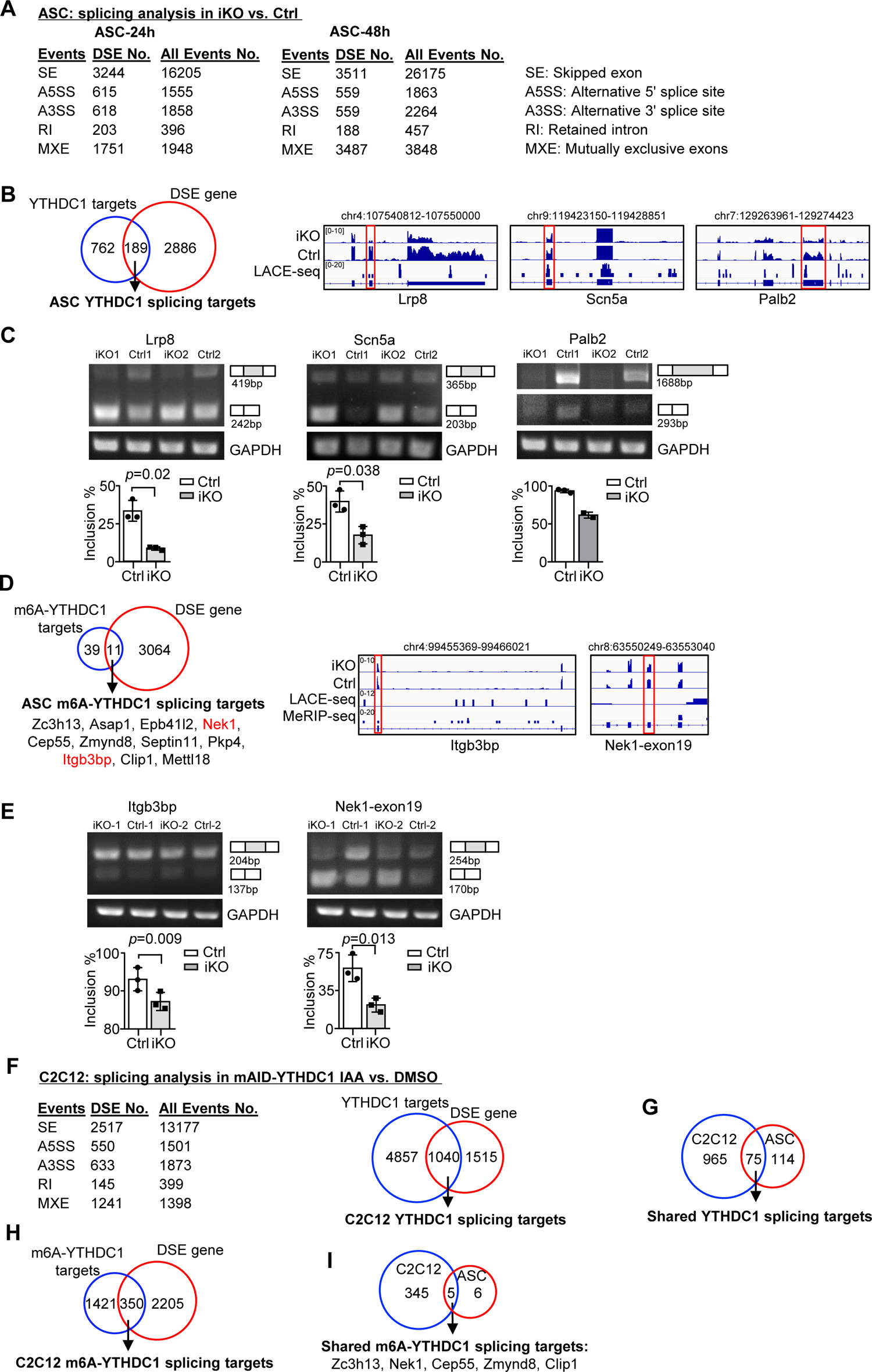
YTHDC1 depletion in ASCs leads to altered splicing events. **A.** Splicing analysis based on bulk RNA-seq data from ASC-24h or −48h defined five types of splicing events. The total number of each event in Ctrl, and the differential spliced events (DSE) in iKO vs. Ctrl are shown. **B.** Left: Combining the above ASC-48h DSEs with the ASC LACE-seq targets identified a total of 189 YTHDC1 splicing target mRNAs. Right: genome tracks of three selected targets. **C**. Top: RT-PCR assay was performed in ASC-48h from YTHDC1-iKO and Ctrl to verify altered splicing of the three selected target mRNAs, Palb2, Lrp8 and Scn5a. GAPDH was used as a control. Bottom: Quantification of exon inclusion level. Exon inclusion level was defined as the percentage of transcripts which includes the specific exon. Included / (Included + Skipped). *n*=3 mice per group for Lrp8 and Scn5a. **D.** Left: combining the above ASC-48h DSE with m6A-YTHDC1 targets uncovered 11 m6A-YTHDC1 splicing targets. Right: genome tracks of two selected targets. **E.** Top: RT-PCR assay was performed in ASC-48 from YTHDC1-iKO and Ctrl to verify altered splicing of the two selected target mRNA, Itgb3bp, and Nek1. GAPDH was used as a control. Bottom: Quantification of exon inclusion level. *n*=3 mice per group. **F.** Left: Splicing analysis based on bulk RNA-seq data from C2C12-mAID-YTHDC1 cells with or without YTHDC1 degradation. Right: Combining the above C2C12 DSEs with the C2C12 LACE-seq targets identified a total of 1040 YTHDC1 splicing target mRNAs. **G.** Overlapping between the above identified YTHDC1 splicing targets in ASC-48h and C2C12. **H.** Combining the above C2C12 DSEs with m6A-YTHDC1 targets uncovered 350 m6A-YTHDC1 splicing targets. **I**. Overlapping of m6A-YTHDC1 splicing targets in C2C12 and ASC-48h. Bars represent mean ± s.d. for all graphs. Statistical significance was determined using a two-tailed Student’s t test.

To ascertain the above findings, we also performed RNA-seq in C2C12 cells with inducible YTHDC1 deletion (cells were treated with IAA for 8h to capture the immediate effect only). A total of 2555 genes with DSE were detected upon YTHDC1 degradation and a large portion (1040) were its binding targets defined in Fig. 4D (Fig. 5F and Suppl. Table S3). This evidence suggests that YTHDC1 is possibly a dominant splicing regulator in C2C12. Of note, these targets were highly enriched for “Chromatin organization”, “Histone modification” (Suppl. Fig. S4B and Suppl. Table S3). 75 of these YTHDC1 splicing targets were also found in ASC (Fig. 5G) and enriched for “Transcription coregulator activity”, “Transcription corepressor activity” (Suppl. Fig. S4C and Suppl. Table S3). Furthermore, 350 out of the 1771 m6A-YTHDC1 targets identified in C2C12 cells (Fig. 4H) were differentially spliced upon YTHDC1 degradation (Fig. 5H) and enriched for “mRNA processing”, “Regulation of mRNA metabolic process” (Suppl. Fig. S4D and Suppl. Table S3); and 5 of these targets were shared in ASCs, including Zch13, Nek1, Cep55, Zmynd6, and Clip1 mRNAs (Fig. 5I). Of note, Nek1 is known to play a role of regulating cell cycle^32^. Altogether, these findings lead us to conclude that m6A-YTHDC1 orchestrate splicing of some mRNAs in myoblast cells.

### YTHDC1 loss inhibits mRNA nuclear export

In addition to splicing regulation, we also wondered if YTHDC1 plays a role in controlling mRNA nuclear export in myoblasts^8, 9^. To test this notion, nuclear (nuc) and cytoplasmic (cyto) fractions were prepared from ASC-48h cells of Ctrl or YTHDC1 iKO muscles and subject to RNA-seq respectively. As a result, a total of 11637 mRNAs were found to be expressed in cyto or nuc fractions and the cyto/nuc ratio was calculated to assess the degree of nuclear export; no significant difference was detected in the average cyto/nuc ratio of these expressed mRNAs in iKO vs. Ctrl (Suppl. Fig. S5A-B and Suppl. Table S4), suggesting YTHDC1 loss may not cause global alteration in mRNA nuclear export. Nevertheless, 1045 mRNAs did show altered cyto/nuc ratio and they were enriched for “Methylation”, “Chromatin modification” etc. (Supp. Fig. S5C and Suppl. Table S4). To specifically examine if YTHDC1 binding could directly modulate nuclear export, we then performed the above analysis on the identified YTHDC1 binding targets (Fig. 4C). A total of 687 were expressed in cyto or nuc fractions and no significant difference was detected in the average cyto/nuc ratio of these mRNAs in the iKO compared to the Ctrl (Fig. 6A-B and Suppl. Table S4), suggesting YTHDC1 binding may not have significant impact on target mRNA nuclear export. Nevertheless, 54 mRNAs did display significant cyto/nuc decrease in iKO vs. Ctrl thus defined as YTHDC1 export targets (Fig. 6C-D and Suppl. Table S4) and they were enriched for “Protein serine kinase activity” (Suppl. Fig. S5G and Suppl. Table S4). Of note, 13 were also defined as YTHDC1 splicing targets (Fig. 6C), suggesting YTHDC1 may simultaneously regulate both splicing and export of these target mRNAs.

**Fig. 6.**
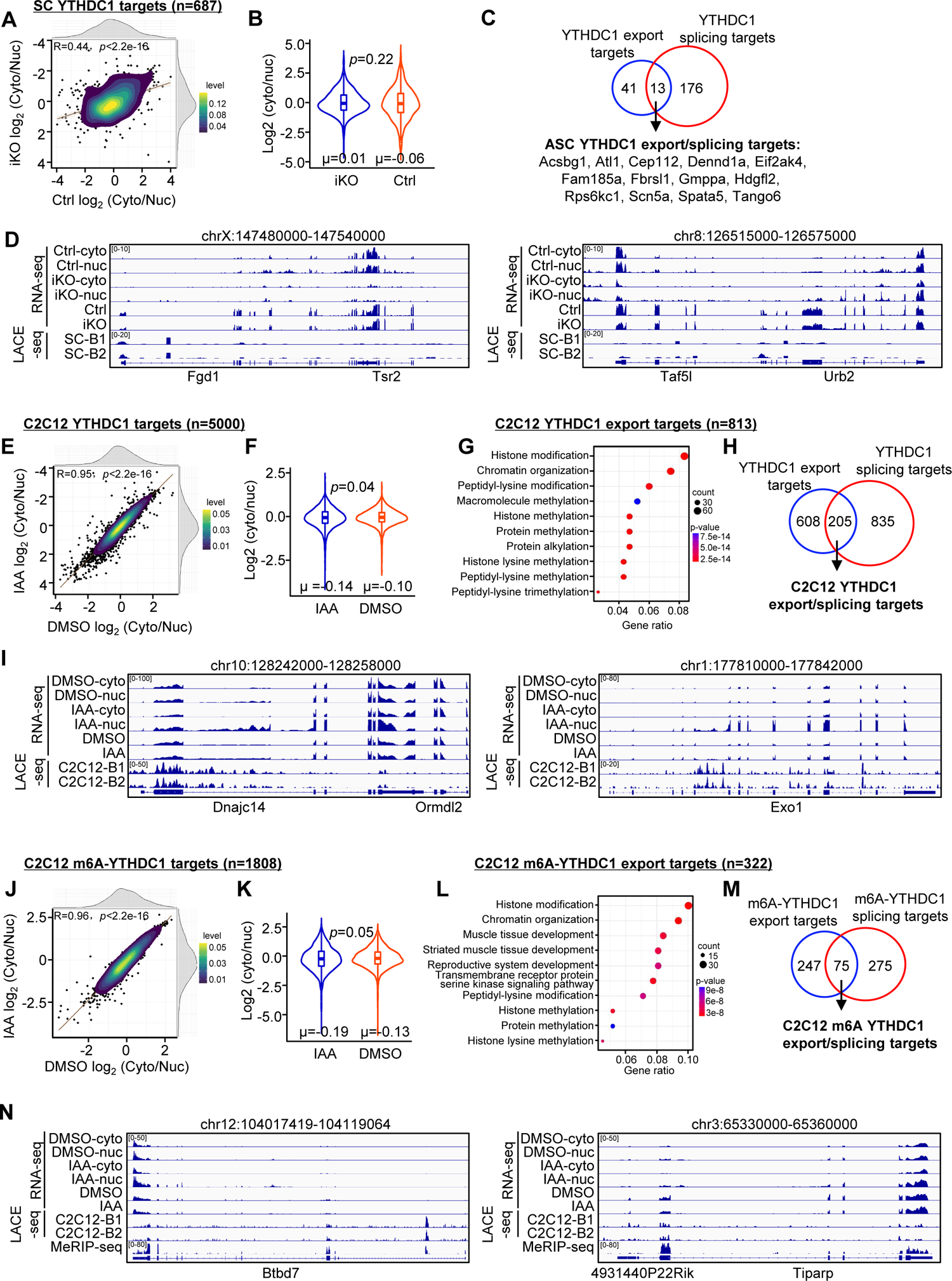
YTHDC1 loss inhibits mRNA nuclear export. **A.** Subcellular RNA-seq was performed using cytoplasmic and nuclear fractions isolated from ASC-48h of Ctrl and YTHDC1 iKO. The log2 (cyto/nuc) expression change was calculated for YTHDC1 targets. On the top and right, the density plot of log2 (cyto/nuc) expression changes is depicted. **B.** Quantification of log2 (cyto/nuc) expression changes in iKO vs. Ctrl. **C.** Overlapping between YTHDC1 mRNA export targets and splicing targets in ASCs. **D.** Genome tracks of two selected export targets. **E-I.** The above assay/analysis was performed in DMSO or IAA treated mAID-YTHDC1 C2C12 myoblasts to identify YTHDC1 regulated export targets in C2C12. **J-N.** The above analyses were conducted using m6A-YTHDC1 targets to identify m6A-YTHDC1 mRNA export targets in C2C12.

We then performed the above assay/analyses in C2C12 mAID-YTHDC1 cells. Similarly, no global impact on nuclear export of all expressed mRNAs (11175) was detected upon YTHDC1 degradation (Supp. Fig. S5D-F and Suppl. Table S4). When analyzing YTHDC1 bound mRNAs, a much higher number (5000) compared to the ASC (because of the higher number of LACE-seq targets from C2C12 (Fig. 4)), were found to be expressed in cyto or nuc fractions and a significant decrease in the average cyto/nuc ratio was found upon YTHDC1 degradation (Fig. 6E-F and Supp. Table S4), suggesting YTHDC1 loss caused significant impact on global mRNA nuclear export in C2C12. Similarly, a total of 813 binding targets were defined as YTHDC1 export targets and displayed reduction in cyto/nuc ratio upon YTHDC1 degradation (Fig. 6G-I and Suppl. Table S4). These targets were enriched for “Histone modification” and “Chromatin organization” etc (Fig. 6G and Suppl. Table S4). And a large portion 205 (∼25%) of them were also identified as YTHDC1 splicing targets (Fig. 6H). Similarly, a total 322 m6A-YTHDC1 export targets were defined in C2C12 cells and also enriched for “Histone modification” and “Chromatin organization” etc (Fig. 6J-N and Suppl. Table S4). And 75 of them were also splicing targets of C2C12 m6A-YTHDC1(Fig. 6M and Suppl. Table S4). Altogether, the above results demonstrate that YTHDC1 binding can indeed promote nuclear export of target mRNAs in myoblast cells.

### Co-IP/MS leads to identification of YTHDC1 interacting partners

Lastly, to further fathom the mechanism underlying the above identified YTHDC1 regulatory functions in myoblasts, we performed Co-immunoprecipitation (Co-IP) coupled with mass spectrometry (MS) to identify its protein interactome knowing that the function of YTHDC1 is largely mediated by its transcriptional or post-transcriptionally interacting partners^12^. To this end, Flag-tagged YTHDC1 or a pRK-vector plasmid was transfected in C2C12 myoblasts (Fig. 7A-B); the interacting proteins were retrieved by anti-Flag beads and subject to MS analysis (Fig. 7A). The results identified a total of 912 potential interacting partners (Fig. 7C and Suppl. Table S5). Of note, they were highly enriched for RNA splicing and mRNA export factors (Fig. 7D), which was in accordance with the above findings, suggesting that the dominant functions of YTHDC1 in myoblasts are splicing and export regulations.

**Fig. 7.**
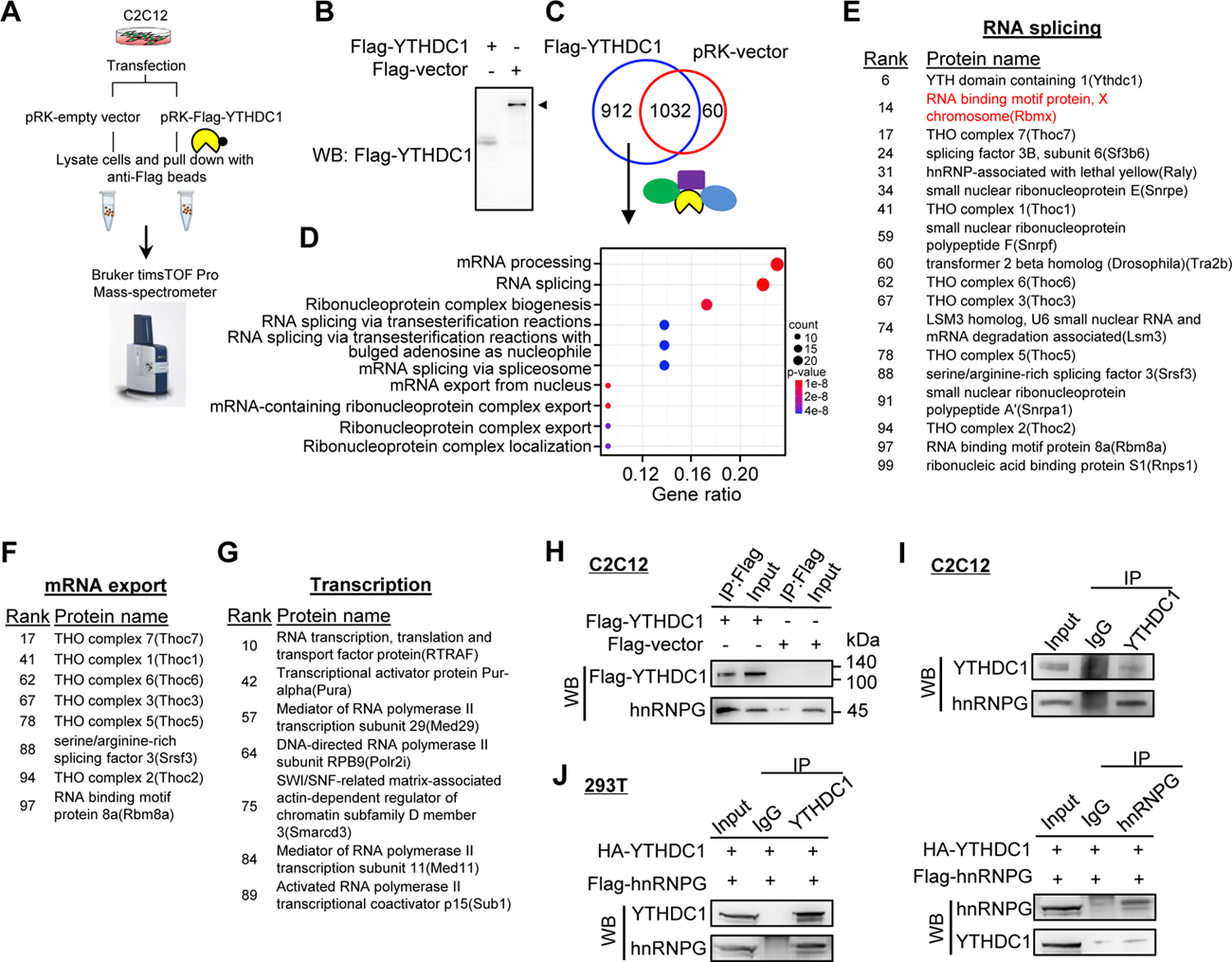
Co-IP/MS leads to identification of YTHDC1 interacting partners. **A-B.** Schematic illustration of the Co-IP/MS procedure. An empty vector or Flag tagged YTHDC1 plasmid was expressed in C2C12 myoblasts followed by pull down with Flag beads; the retrieved proteins were subject to MS with Bruker timsTOF Pro. **B.** Overexpression of the Flag tagged YTHDC1 was confirmed by WB using anti-Flag antibody. **C.** 912 proteins were uniquely retrieved in YTHDC1 but not Vector expressing cells. **D.** GO functions of the above proteins are shown. **E-G.** The above identified interacting proteins with RNA splicing, mRNA export, or potential transcriptional regulatory functions are shown in the lists. **H.** Flag tagged YTHDC1 was overexpressed in C2C12 and Flag-beads based IP was performed followed by WB to verify the retrieved hnRNPG protein. **I.** IP of endogenous YTHDC1 protein in C2C12 myoblasts followed by WB to examine retrieved hnRNPG protein. **J.** Flag-hnRNPG and HA-YTHDC were overexpressed in 293T cells; IP with YTHDC1 or hnRNPG protein flowed by WB to confirm the interaction between the two proteins.

Among the splicing factors (Fig. 7E), hnRNPG (also called RNA binding motif protein X, RBMX) ranked very high (No. 14) on the list. It is a ubiquitously expressed RBP best known for its function in splicing control via interactions with SRSF3, SRSF7, SLM, SAFB1^33, 34^; interestingly, hnRNPG was recently defined as an m6A reader protein and it can regulate splicing through simultaneous interacting with m6A modified nascent transcripts and RNAPII^35^. hnRNPG thus represents a versatile RBP like YTHDC1 in both splicing and transcriptional regulations. Their physical interaction was validated by performing Co-IP followed by WB using both exogenously (Fig. 7H) or endogenously (Fig. 7I) expressed YTHDC1 in C2C12 myoblasts; additionally, the interaction was also validated in 293T cells (Fig. 7J). Furthermore, we also identified the well-known binding partners of YTHDC1, SRSF3^6^ on the list (Fig. 7E), strengthening its function in regulating splicing in myoblasts as characterized in Fig. 5.

Examining the list of mRNA export factors (Fig. 7F), several components of the THO subcomplex of the TREX (Transcription Export) complex were identified. The bulk of mRNA nuclear export is a complex yet well characterized process mediated by the TREX complex and the heterodimeric nuclear export receptor NXF1:NXT1^8, 36^. As the key initiating complex, TREX has 14 known subunits with the multimeric THO complex sitting as the core. Strikingly, all the known THOC members including THOC7, 1, 6, 3, 5, and 2 were retrieved, reinforcing YTHDC1 function in controlling mRNA export as defined in Fig. 6. Furthermore, we also identified some known transcriptional regulators including RTRAF, Med 29 and Med 11 (Fig. 7G), pointing to previously unknown means via which YTHDC1 may regulate transcription. Altogether, the results from the Co-IP/MS are in line with findings from Figs. 5 and 6 to reinforce that the notion that YTHDC1 play pleiotropic regulatory functions in myoblasts.

## Discussion

In this study we investigate the functional role of m6A reader YTHDC1 protein in skeletal muscle stem cells and muscle regeneration. Our findings demonstrate the expression dynamics of several m6A regulators including writers, readers and erasers during the course of SC lineage progression, implicating their possible involvement in governing SC activities. Among these m6A regulators, we characterize YTHDC1 function in depth and uncover it as an essential factor controlling SC activation and proliferation. Inducible depletion of YTHDC1 in SCs drastically impairs SC activation and proliferation thus almost abolishes acute injury induced muscle regeneration. Mechanistically, LACE-seq identifies YTHDC1 binding mRNA targets among which a portion are m6A dependent. Further splicing analyses provide evidence for m6A-YTHDC1 participation in modulating splicing events in myoblast cells. Additionally, subcellular fractionation RNA-seq also defines potential mRNA export targets of YTHDC1 in myoblasts. Lastly, Co-IP/MS defines a wide array of interacting protein partners of YTHDC1 including mRNA splicing and export factors as well as transcriptional regulators, and they may function in synergism to mediate the pleiotropic functions of YTHDC1 in myoblasts. Among these factors, hnRNPG appears to be a bona fide functional partner of YTHDC1. Altogether, our findings demonstrate that YTHDC1 is an indispensable intrinsic regulator of SC activities and muscle regeneration through multi-faceted controlling of RNA metabolism in myoblast cells (Fig. 8).

**Fig. 8.**
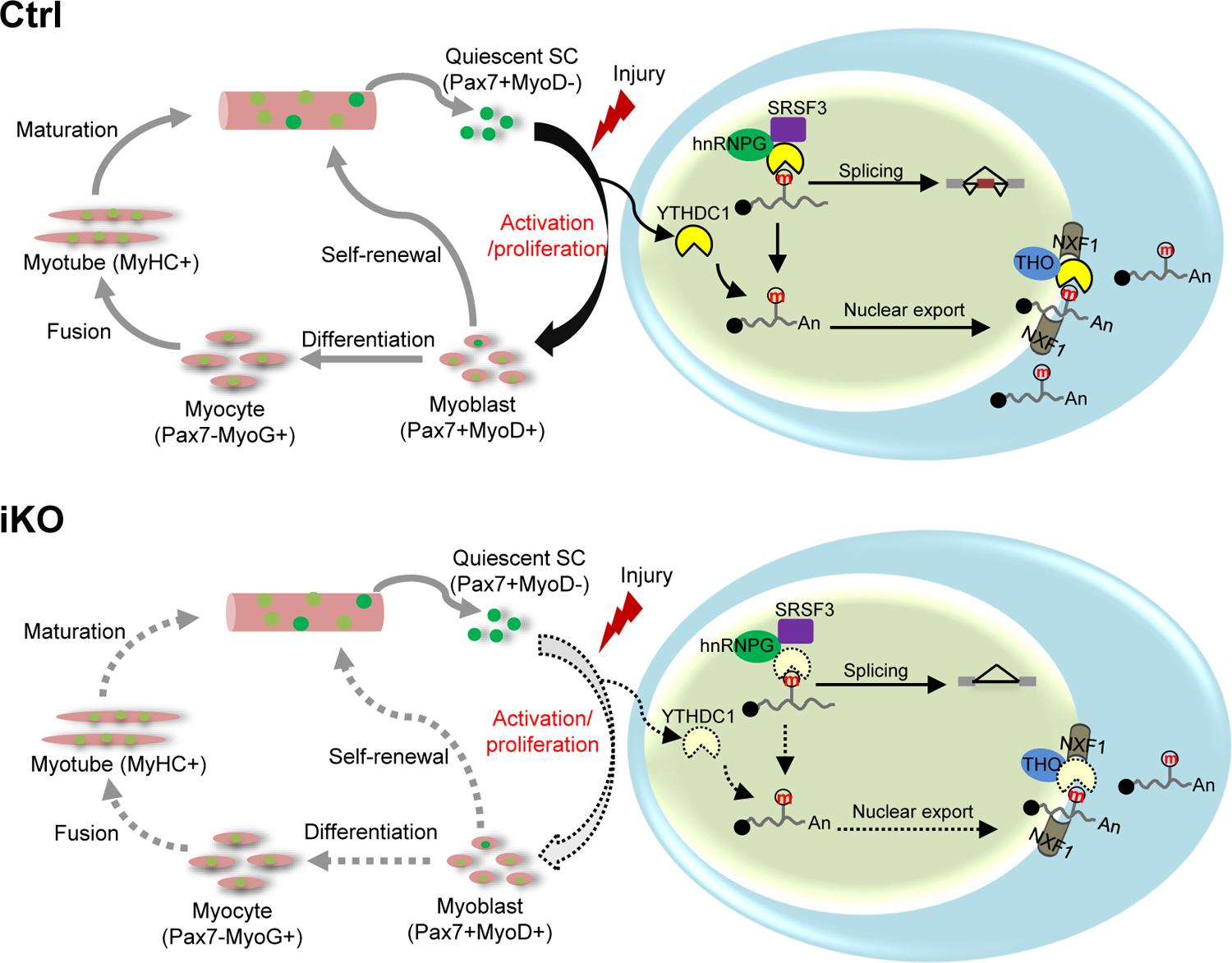
A working model of YTHDC1 function in SCs. YTHDC1 is induced upon SC activation and proliferation to promote SC activation and proliferation upon acute injury induced muscle regeneration. Mechanistically, it functions through both facilitating mRNA splicing synergistically with hnRNPG and promoting mRNA export possibly by binding with the THO nuclear export complex.

Understanding the intrinsic gene regulatory mechanisms governing SC activities is no doubt the foundation to decipher SC contribution to muscle regeneration. Compared to the wealth of knowledge in transcriptional regulation, post-transcriptional regulation in SCs remains largely unexplored but emerging evidence supports its importance. For example, some mRNAs are transcribed and stored in the quiescent SCs to allow the rapid production of protein products upon SC activation^37–40^. Our recent findings also demonstrate that a RNA helicase, DHX36 is an indispensable post-transcriptional regulator of skeletal muscle regeneration through diverse modes of targeting and regulation^3^. Nevertheless, until now the study of m6A regulation and function in SCs is scarce ^21–23^. Our current study is the first to identify potential functional m6A regulators and provides a holistic investigation of a writer protein, YTHDC1 function in SCs and muscle regeneration. The transcriptomic profiling uncovered that an array of m6A regulators are dynamically expressed in the course of SC lineage progression; notably, YTH-domain containing reader proteins including YTHDC1, YTHDF1, YTHDF2 and YTHDF3 were all highly expressed in SCs. The mRNA level of YTHDC1 was high across the entire course albeit showing an induction in FISC; its protein level, nevertheless, was induced in ASC-24h and continued to increase until ASC-48h, suggesting its dominant role in governing activation/proliferation. Indeed, the subsequent dissection using the YTHDC1-iKO mouse provided solid evidence supporting its positive role in promoting SC activation and proliferation; inducible depletion of YTHDC1 largely inhibited SC activation and proliferation (Fig.2). We believe the incompetence of SC activation is the major cause of nearly blocked regeneration after acute injury induced muscle damage. Nevertheless, the potential defects in other aspects such as differentiation, self-renewal and survival need to be investigated in the future. Our findings thus suggest that YTHDC1 plays an indispensable role in SCs and muscle regeneration, which is not replaceable by other m6A readers. This study also adds genetic evidence for the importance of YTHDC1 in cellular processes, which is largely lacking in the field. In the future, it will also be interesting to test the functionality of other readers such as YTHDF1, 2 and 3 in SCs.

To investigate the functional mechanism of YTHDC1 in promoting SC activation/proliferation, recently developed LACE-seq was harnessed using both ASCs and C2C12 myoblasts. Even though the number of peaks identified in ASCs was not comparable with the number in C2C12, LACE-seq has enabled the profiling of transcriptome-wide YTHDC1 binding in ASCs, which was previously impossible using traditional mRNA binding probing methods. By combining with the m6A MeRIP-seq dataset available in C2C12^31^, we were able to define m6A modified binding targets of YTHDC1. Of note, a large portion of the YTHDC1 binding targets do not seem to possess m6A modification (Fig. 4H), suggesting YTHDC1 may have m6A independent functions. In the future, it will be necessary to generate transcriptome-wide m6A mapping in SCs, which has become possible by the recently developed methods enabling the use of low number of cells. Nevertheless, with the available datasets, we were able to define sets of m6A-YTHDC1 mRNA targets in myoblasts and these are interestingly enriched for RNA splicing/processing genes.

Next, we examined YTHDC1 regulation of mRNA splicing in depth by examining altered splicing events upon YTHDC1 loss, which unveiled the splicing targets of m6A-YTHDC1. The number is relatively small in ASCs due to the relatively small number of YTHDC1 binding targets from LACE-seq; a much higher number was found in C2C12 cells but constitutes only a small number (20%) of m6A-YTHDC1 targets. Additionally, we examined potential effect of YTHDC1 loss in mRNA nuclear export and indeed identified a set of mRNAs possibly regulated by m6A-YTHDC1; upon YTHDC1 loss, the nuclear enrichment level was significantly increased. Interestingly, a large percentage of these mRNAs are also splicing targets of m6A-YTHDC1, suggesting YTHDC1 binding to a m6A modified mRNA can control mRNA metabolism at multiple levels. Of note, our study identifies a large number of splicing or/and export targets of m6A-YTHDC1 and they play a wide range of cellular functions directly or indirectly related to cellular proliferation. The observed proliferative defects upon YTHDC1 loss thus could be a synergistic effect of many down-stream targets.

Lastly, Co-IP/MS was performed in myoblasts and led to the identification of an array of RNA processing regulators mainly including splicing and nuclear export factors. This finding is in concordance with the above demonstrated YTHDC1 functions in splicing and export regulations. In addition, transcriptional regulators were also identified. The m6A dependent functions of YTHDC1 in regulating splicing, export or transcription have all been individually reported in various cell contexts^7, 9, 19^, our finding thus hints pleiotropic roles of YTHDC1 in myoblasts. In terms of splicing regulation, Xiao W et. al. showed that YTHDC1 binds competitively to the splicing factor SRSF3, while antagonizing the binding of SRSF10 to promote exon inclusion^6^. The top ranked binding partner on our list, however, was not SR splicing factors (Fig. 7E), instead, hnRNPG appeared as the most likely interacting partner. Its specific interaction with YTHDC1 was validated by Co-IP/WB using both exogenously and endogenously expressed proteins in C2C12 and also in 293T cells (Fig. 7), thus pointing to hnRNPG as a bona fide interacting partner of YTHDC1. Interestingly, hnRNPG also represents a multi-faceted gene regulator. Originally known as a hnRNP family splicing factor via interactions with SRSF3, SRSF7, SLM, SAFB1^33, 34^, it was later shown to bind with m6A modified mRNAs thus possessing m6A reader function. It can co-transcriptionally interact with both RNAPII and m6A-modified nascent pre-mRNA to modulate RNAPII occupancy and alterative splicing^35^. Its implicated role in transcription repression was also reported^33^. Therefore both hnRNPG and YTHDC1 are versatile m6A dependent gene regulators yet no study so far has made functional connection between these two key m6A readers despite their physical interaction was mentioned^20, 34^. The future efforts will be focused on further delineating the regulatory and functional synergism between YTHDC1 and hnRNPG in SCs. In addition, the Co-IP/MS also identifies a number of TREX components as potential interacting partners of YTHDC1, and a prior report also demonstrated the physical interaction between TREX complex with YTHDC1 in HEK293T cells^8^. The modulation of nuclear export by YTHDC1 is thus very likely exerted together with TREX proteins, which will need to be further investigated in the future.

Altogether, biologically, our findings uncover YTHDC1 as an essential post-transcriptional regulator of SC activities and muscle regeneration. In the coming years, more solid genetic evidence will be needed to demonstrate the key roles of m6A regulators in various biological systems. It will also be interestingly to determine if YTHDC1 deregulation is implicated in muscle related diseases such as aging associated sarcopenia. Mechanistically, we demonstrate multi-faceted roles of YTHDC1 in decoding m6A-marked transcripts in myoblasts; a wide range of mRNA targets are controlled by YTHDC1 thus mediating its effect in promoting myoblast proliferation. Although our current study mainly focuses on delineating it post-transcriptional actions, we believe YTHDC1 also modulate transcriptional events in myoblast nucleus and hnRNPG seems to constitute an important co-factor that may synergistically regulate both splicing and transcription functions of YTHDC1. In addition, it will be interesting to explore if YTHDC1, hnRNPG, and the target mRNAs self-organize into distinct condensates within the myoblast nucleus as emerging evidence suggests RNA, m6A readers and co-factors can cooperatively form distinct nuclear condensates at specific genomic loci^12^.

## Methods

### Mice

*Pax7^CreER^ (Pax7^tm1(cre/ERT2)Gaka^)* and Tg: Pax7-nGFP mouse strains were kindly provided by Dr. Shahragim Tajbakhsh. *Ythdc1^fl^(B6;129S4-Ythdc1tm1.1Jw/Mmjax)* strain and *ROSA^EYFP^* mouse were provided by Jackson Laboratory. *Pax7^CreER^* mice were crossed with *ROSA^EYFP^*mice to generate the *Pax7^CreER^; ROSA^EYFP^* reporter mice. The *Ythdc1* inducible knock out (*Ythdc1* iKO) mice with EYFP reporter (Ctrl: *Pax7^CreER/+^; ROSA^EYFP/+^; Ythdc1^+/+^*, iKO: *Pax7^CreER/+^; ROSA^EYFP/+^; Ythdc1^fl/fl^*) were generated by crossing *Ythdc1^fl^* mice with *Pax7^CreER^; ROSA^EYFP^* reporter mice. The mice were maintained in animal room with 12h light/12 h dark cycles, 22–24 °C room temperature and 40–60% humidity at the animal facility in the Chinese University of Hong Kong (CUHK). All animal handling procedures and protocols were approved by the Animal Experimentation Ethics Committee (AEEC) of CUHK (Ref. No. 19-251-MIS). All animal experiments with iKO mice followed the regulations and guidance of laboratory animals set in CUHK.

### Animal procedures

Inducible conditional deletion of *Ythdc1* was administered by injecting Tamoxifen (Tmx) (T5648, Sigma) intraperitoneally (IP) at 2mg per 20 g body weight. For BaCl_2_ induced muscle injury, 2-3-month-old mice were intramuscularly injected with 50 μl of 10 μg/ml BaCl_2_ solution into TA muscles and the muscles were harvested at designated time points for further analysis. For EdU incorporation assay *in vivo*, 2 days after BaCl_2_ injection, EdU was injected via IP at 0.25 mg per 20 g body weight, followed by FACS isolation of SCs 12h later. SCs were then seeded and fixed with 4% PFA for further stain and analysis.

### Satellite cell isolation by FACS

Briefly, hindlimb muscles from *Ythdc1* Ctrl/iKO and Pax7n-GFP mice were dissected and minced with blades, then digested with collagenase II (1100 U ml^-1^, Worthington in Hams F-10 media (Sigma)) for 90 min at 37°C with gentle rotation at 70 rpm. The digested muscles were washed in washing medium (Hams F-10 media, 10% HIHS (Gibco), penicillin/streptomycin (1×, Gibco) once and SCs were further released by treating muscles with Collagenase II (1100 U ml^-1^) and Dispase (11 U ml^-1^) for 40 min at 37°C. Digested tissue was passed through a 21-gauge needle 12 times and filtered through a 40-μm filter followed by spinning at 700 × g for 5 min at 4 °C. Mononuclear cells were resuspended and filtered with a 40-µm cell strainer and GFP+/EYFP+ SCs were sorted out by BD FACS Aria Fusion cell sorter (BD Biosciences).

### Single myofiber isolation

Single myofibers were isolated as previous described^3^. 2-3-month-old mice were used for extensor digitorum longus muscles (EDL) isolation. After incubation in collagenase II solution (800U/ml) at 37 °C in a water bath for 70 min, muscles were transferred to a dish containing pre-warmed washing medium (F10+10% Horse serum+ Penicillin-Streptomycin). Single myofibers were released by pipetting with a large hole bore glass pipette gently and transferred to a new dish with culture medium (F10+10% Horse serum+ Penicillin-Streptomycin+ b-FGF (0.025μg/ml)) for *ex-vivo* culture. Fibers were fixed with 4% PFA at designed time point. For EdU assay, 10μM EdU was added to culture medium for 4hr before fixation.

### Cell culture

Mouse C2C12 myoblast cells (CRL-1772) and 293T cells (CRL-3216) were obtained from American Type Culture Collection (ATCC) and cultured in DMEM medium with 10% fetal bovine serum, 100 units/ml of penicillin, and 100 μg of streptomycin (growth medium, or GM) at 37 °C in 5% CO2. Freshly isolated SCs were cultured in Ham’s F10 medium supplemented with 20% FBS and bFGF (0.025 μg/ml) (growth medium) on dish/slides precoated with PDL and ECM, ASC-24h and ASC-48h were harvested at designed time point for RNA/protein/Immunofluorescence analysis.

### Generation of mAID-YTHDC1 inducible degradation C2C12 cells

Briefly,^41^ OsTIR1 (F74G) mutant was first introduced into PB-Ostir-neo(Addgene plasmid #161973) plasmid by overlap PCR to generate AID2 system in C2C12 cells with improved degradation efficiency and reduced background degradation. CRISPR–Cas9 plasmid was generated using PX458 (Addgene plasmid #48138) and sgRNA targeting N-terminal of YTHDC1 near ATG start codon. Two donor plasmids were generated by overlap PCR ∼500bp *Ythdc1* genomic sequence flanking BSD/HygR-P2A-mAID-mCherry2 sequence (Addgene plasmid #121180, #121183) and cloning into pMD20-T vectors by T-A ligation. Cas9 and donor plasmids were transfected to C2C12 (Ostir2) cells and antibiotic selection (BSD 10ug/ml, Hygro 100ug/ml) was performed for 1 week before single clone selection. Successful knocked in clones were validated by genotyping PCR and Western blot. 1μM 5-Ph-IAA was used to induce mAID-YTHDC1 degradation with DMSO treatment used as a control.

### EdU Assay

EdU assay was performed following manufacture’s protocol (Invitrogen™, Click-iT™ EdU Cell Proliferation Kit for Imaging, Alexa Fluor™ 594 dye, C10339). Cells/myofibers were incubated with 10μM EdU for 4hr before fixation with 4% PFA.

### Plasmids

pRK5-flag-YTHDC1 and pRK5-flag hnRNPG plasmids were generated by amplifying ORF of YTHDC1 and hnRNPG from SCs cDNA and cloned into pRK5-flag empty vector through NotI and HindIII restriction sites.

### Cell fractionation

Cell fractionation protocol was modified based on previous protocol^42^. C2C12 or SCs were collected in cold PBS, washed once and then incubated in buffer A (HEPES-KOH 50 mM pH 7.5, 10 mM KCl, 350 mM sucrose, 1 mM EDTA, 1 mM DTT, 0.1% Triton X-100) for 5 min on ice and then homogenized with a T 10 basic ULTRA-TURRAX® homogenizer at 4^th^ gear for 1min. The nuclei were harvested by brief centrifugation (2000g, 5 min), while the supernatant was collected as the cytoplasmic fraction. Nuclei were resuspended with same volume buffer A as supernatant. RNAs were extracted using TRIzol reagent and further analyzed by RT-qPCR and RNA-seq. For proteins, 6xSDS loading buffer were added accordingly for western blot analysis.

### RNA isolation, and quantitative RT-PCR

**T**otal RNAs were extracted using TRIzol reagent (Invitogen) follow manufacture’s protocol. For quantitative RT-PCR, cDNAs were revers transcribed using HiScript III 1^st^ Strand cDNA Synthesis Kit (Vazyme, R312-01). Real-time PCR reactions were performed on a LightCycler® 480 Instrument II (Roche Life Science) using Luna Universal qPCR Master Mix (NEB, M3003L). For splicing verification, GAPDH was used as control. cDNA of Ctrl/iKO were amplified by 30 cycles of PCR and run on 2% agarose gel. Sequences of all primers used can be found in Suppl. Table S7.

### RNA-seq and data analysis

For RNA-seq (polyA+ mRNA), total RNAs were subject to polyA selection (Ambion, 61006) followed by library preparation using NEBNext® Ultra™ II RNA Library Preparation Kit (NEB, E7770S). Libraries were paired-end sequenced with read lengths of 150 bp on Illumina HiSeq X Ten or Nova-seq instruments. The raw reads of RNA-seq were processed following the procedures described in our previous publication. Briefly, the adapter and low-quality sequences were trimmed from 3’ to 5’ ends for each read, and the reads shorter than 50 bp were discarded. The clean reads were aligned to mouse (mm9) reference genome with STAR. Next, we used Cufflinks to quantify the gene expression. Genes were identified as differentially expressed genes (DEGs) if the change of expression level is greater than 2 folds and the p-value is less than 0.01 between two stages/conditions. GO enrichment analysis were performed using R package clusterProfiler.

### Immunoblotting and immunofluorescence

Immunoblotting and immunofluorescence were performed according to our standard protocols^3, 43–45^. Proteins were extracted using RIPA lysis buffer. The following dilutions of antibodies were used for western blot staining: PAX7 (Developmental Studies Hybridoma Bank; 1:200), GAPDH (Sigma, G9545, 1:3000), YTHDC1 (Cell Signaling Technology, #77422, 1:1000), MyoD (Abclonal, 1:1000), hnRNPG (Cell Signaling Technology, #14794, 1:1000), Histone H3 (Santa Cruz, sc-517576, 1:5000), Flag-tag (Sigma, F3165, 1:3000). For immunofluorescence staining, cultured cells and myofibers were fixed with 4% PFA at RT for 15min, after washing with PBS, cells were permeabilized with 0.2% TritonX-100 and followed by 30min blocking with 3% BSA. Primary antibodies were incubated at 4℃ overnight. Following antibodies and related dilutions were used: PAX7 (Developmental Studies Hybridoma Bank; 1:50), eMyHC (Leica NCL-MHC-d;1:200), laminin (Sigma-Aldrich L9393; 1:800), MyoD (Abclonal, 1:1000), YTHDC1 (Abcam, ab122340, 1:200). Secondary antibodies were incubated at RT for 1h and nuclei were stained with ProLong™ Gold Antifade Mountant with DAPI (Thermo). Immunofluorescence staining of frozen muscle sections was performed as previously described^43^. Briefly, samples were boiled in 0.01M citric acid (pH 6.0) for 10 min in a microwave before blocking with 4% BBBSA (4% IgG-free BSA in PBS; Jackson, ref: 001-000-162). Then the endogenous mouse IgG were blocked by treatment with the Donkey anti-Mouse IgG (H+L) (1/100 in PBS; Jackson, ref: 115-007-003) for 30 min. After primary antibody incubation overnight, the biotin conjugated anti-mouse IgG (1:500 in 4% BBBSA, Jackson, ref: 205) and Cy3-Streptavidin (1:1250 in 4%BBBSA, Jackson, ref: 016–160) were used as secondary antibodies for Pax7 staining. All fluorescent images were captured with a fluorescence microscope (Leica DM6000 B). H&E staining on frozen muscle sections was performed as previously described^45^.

### Co-IP/Mass spectrometry

Immunoprecipitation of C2C12 cell overexpressing flag-YTHDC1 was performed based on previous publication^44^. Briefly, C2C12 cells were transfected with pRK5-flag-YTHDC1 or pRK5-falg empty vector for 48hrs. Cells were digested with trypsin and washed with PBS once, then lysed in hypotonic lysis buffer (10mM HEPES, pH 7.9, 10mM KCl, 0.1mM EDTA, 0.1mM EGTA, and complete protease inhibitors (Roche)) and incubated on ice for 15 min. The lysates were centrifuged for 10 min at 2000 g and supernatant was discarded. The pelleted nuclei were washed once with hypotonic lysis buffer and then resuspended in hypertonic buffer (20mM HEPES, pH 7.9, 0.4M NaCl, 1mM EDTA, 1mM EGTA, 0.6% NP-40 and complete protease inhibitors (Roche)), digested with the DNase I (AM2238, Thermo Fisher Scientific) for 45min at 4 °C, and spun down at 12000g for 10min at 4 °C. The nuclear lysates were diluted 2 fold with IP buffer (20mM HEPES, pH 7.9, 0.2M NaCl, complete protease inhibitors (Roche) and then flag-beads (Anti-Flag Magnetic Beads, MCE, HY-K0207) were added and incubated overnight at 4 °C. After 5 time of washing, proteins were eluted with elution buffer (62.5mM Tris-HCL 7.5, 0.2% SDS) at 99°C for 10min, 20 percent elution was subjected to western blot verification. Mass spectrometry experiment was performed using the Bruker timsTOF Pro Mass-spectrometer with the help of Biosciences Central Research Facility, HKUST. Mass spectrometry raw data was processed by PEAKS software (Version: X +). Protein abundance was obtained by normalizing the spectral number of proteins with the length of the protein. Unique YTHDC1 interacting proteins were selected comparing flag-YTHDC1 sample and flag-vector sample (excluding nonspecific binding targets).

### Co-IP

Co-IP of C2C12 cells and 293T cells was performed based on published protocol^46^. Briefly, cells were incubated in buffer A for 5 min on ice and then homogenized as described for cell fractionation. The nuclei were pelleted by 2000g, 5 min centrifugation and resuspended in hypertonic buffer (20 mM HEPES, pH 7.5, 10% glycerol, 0.42 M KCl, 4 mM MgCl2, 0.2 mM EDTA, 0.5 mM DTT, Protease Inhibitor Cocktail). After 30 min incubation on ice, nuclear extract was collected by high-speed centrifugation (12,000 rpm, 15 min, 4 °C). Same volume of hypotonic buffer (10 mM HEPES pH 7.5, 1.5 mM MgCl2, 10 mM KCl, 0.5 mM DTT, 1× Protease Inhibitor cocktail) was added to nuclear extract. Lysate was precleared with protein G beads (Dynabeads™ Protein G for Immunoprecipitation, Invitrogen™, 10003D) at 4 °C for one hour with rotation and 10% was saved as Input. After incubation with indicated antibodies overnight, protein G beads were added and incubated for another 2 hours at 4 °C. Beads were washed 5 times with IP buffer and proteins were eluted with 1xSDS loading buffer at 99°C for 5min.

### LACE-seq and data analysis

LACE-seq was performed as described.^29^ Cells were washed with ice cold PBS and subjected to 400mJ UV treatment to crosslink proteins with their interacting RNAs. After crosslink, cells were pelleted and kept at −80°C before library preparation. YTHDC1 antibody (Abcam, ab122340) was used to pull down specific protein-RNA complex from lysate. In brief, MNase was used to cut RNAs into YTHDC1-associated short fragments on beads. The 3’ ends of fragmented RNAs were then dephosphorylated and ligated with a 5’ pre-adenylated linker containing four randomized nucleotides. Biotinylated primer containing the T7 promoter was used for reverse transcription which stopped at crosslinking site (YTHDC1 binding site). cDNAs were poly(A) tailed and purified with streptavidin beads. Second-strand cDNAs were synthesized on beads with an adaptor containing oligo-(dT). After a pre-amplifying step by PCR, T7 RNA polymerase was used to amplify trace amounts of truncated cDNAs linearly. Then the products were PCR converted into libraries for single end sequencing on Illumina platform (Nextseq 550). For the LACE-seq data analysis, first, the adapter sequences and poly(A) tails at the 3’ end of raw reads were removed using Cutadapt (v.1.15) with following parameters: -f fastq -m 18 -n 2 -a A{15} --quality-base=33. Clean reads were first aligned to mouse pre-rRNA using Bowtie software (v.1.2.3) with default parameters, and the remaining unmapped reads were then aligned to the mouse (mm9) reference genome with Bowtie parameters: --best –strata -v 2 -k 10. Pearson’s correlation coefficient between LACE-seq replicates was performed using multiBamSummary module of deeptools. LACE-seq peaks are called by Piranha software (http://smithlabresearch.org/software/piranha/, v.1.2.1) in ASC cells. The parameters were as follows: -sort -p_threshold 0.001 -b 20 -d ZeroTruncatedNegativeBinomial. For motif analysis, LACE-seq peaks were first extended 20 nt to upstream and downstream, respectively. Enriched motifs are scanned by findMotifsGenome.pl from Homer.

### Whole cell RNA-seq and RNA splicing analysis

For the splicing analysis, RNA-seq reads were first aligned to the mm9 reference genome using STAR, splicing events were then detected by rMATS with default parameters^47^. A cutoff of absolute value of “IncLevelDifference” <0.1 was used to define differentially splicing events (DSE) in YTHDC1 iKO vs. Ctrl. DSE targets are the genes that having DSE in its transcripts.

### Subcellular RNA-seq and mRNA exporting analysis

For the subcellular RNA-seq analysis, reads were aligned to reference genome and the quantification of gene expression was performed by Cufflink. A cutoff of FPKM>1 either in cytoplasmic (cyto) or nuclear (nuc) was used to determine the expression of genes. To identify the nuclear enriched mRNAs upon YTHDC1 knockout or degradation, the log_2_(cyto/nuc) of knockout or degradation should be less than the ctrl sample and the log_2_(cyto/nuc) of knockout or degradation should be less than −1.

## Supporting information

supplemental infomation

## Acknowledgments

We thank Dr. Lifang Han in assisting us performing the Mass Spectrometry at Biosciences Central Research Facility, HKUST. This work was supported by the National Natural Science Foundation of China (NSFC) to H.W. (project code: 31871304 and 82172436); General Research Funds (GRF) from the Research Grants Council (RGC) of the Hong Kong Special Administrative Region(14100018, 14115319, 14100620, and 14106521 to H.W.; 14120420, 14116918 and 14120619 to H.S.); the research funds from Health@InnoHK program launched by Innovation Technology Commission, the Government of the Hong Kong SAR, China to H.W.;NSFC/RGC Joint Research Scheme (project code: N_CUHK 413/18 to H.S.); Collaborative Research Fund (CRF) from RGC to H.W. (project number: C6018-19GF); Theme-based Research Scheme (TRS) from RGC (projectnumber:T13-602/21-N); Area of Excellence Scheme (AoE) from RGC(project number: AoE/M-402/20); Natural Science Foundation of Guangdong Province to X.C. (project code: 2019A1515010670).

## Author Contributions

Y.Q. designed and performed most experiments; X.C performed YTHDF1 and YTHDF2 western blot in ASCs and helped with some mouse related experiments; D.W. and R.S. performed LACE-seq of YTHDC1 in C2C12 and ASCs; Q.S. performed all bioinformatics analyses; H.S. supervised computational analyses; Y.Q., Q.S., and H.W. wrote the manuscript, with input from all authors.

## Reference

1. Fujita, R. & Crist, C. Translational Control of the Myogenic Program in Developing, Regenerating, and Diseased Skeletal Muscle. Curr Top Dev Biol 126, 67–98 (2018).

2. Relaix, F. et al. Perspectives on skeletal muscle stem cells. Nat Commun 12, 692 (2021).

3. Chen, X. et al. Translational control by DHX36 binding to 5’UTR G-quadruplex is essential for muscle stem-cell regenerative functions. Nat Commun 12, 5043 (2021).

4. Shi, H., Wei, J. & He, C. Where, When, and How: Context-Dependent Functions of RNA Methylation Writers, Readers, and Erasers. Mol Cell 74, 640–650 (2019).

5. Murakami, S. & Jaffrey, S.R. Hidden codes in mRNA: Control of gene expression by m(6)A. Mol Cell 82, 2236–2251 (2022).

6. Xiao, W. et al. Nuclear m(6)A Reader YTHDC1 Regulates mRNA Splicing. Mol Cell 61, 507–519 (2016).

7. Kasowitz, S.D. et al. Nuclear m6A reader YTHDC1 regulates alternative polyadenylation and splicing during mouse oocyte development. PLoS Genet 14, e1007412 (2018).

8. Lesbirel, S. et al. The m(6)A-methylase complex recruits TREX and regulates mRNA export. Sci Rep 8, 13827 (2018).

9. Roundtree, I.A. et al. YTHDC1 mediates nuclear export of N(6)-methyladenosine methylated mRNAs. Elife 6 (2017).

10. Shima, H. et al. S-Adenosylmethionine Synthesis Is Regulated by Selective N(6)-Adenosine Methylation and mRNA Degradation Involving METTL16 and YTHDC1. Cell Rep 21, 3354–3363 (2017).

11. Zhang, Z. et al. YTHDC1 mitigates ischemic stroke by promoting Akt phosphorylation through destabilizing PTEN mRNA. Cell Death Dis 11, 977 (2020).

12. Widagdo, J., Anggono, V. & Wong, J.J. The multifaceted effects of YTHDC1-mediated nuclear m(6)A recognition. Trends Genet 38, 325–332 (2022).

13. Kan, R.L., Chen, J. & Sallam, T. Crosstalk between epitranscriptomic and epigenetic mechanisms in gene regulation. Trends Genet (2021).

14. Wei, J. & He, C. Chromatin and transcriptional regulation by reversible RNA methylation. Curr Opin Cell Biol 70, 109–115 (2021).

15. Liu, J. et al. The RNA m(6)A reader YTHDC1 silences retrotransposons and guards ES cell identity. Nature 591, 322–326 (2021).

16. Chen, C. et al. Nuclear m(6)A reader YTHDC1 regulates the scaffold function of LINE1 RNA in mouse ESCs and early embryos. Protein Cell 12, 455–474 (2021).

17. Li, Y. et al. N(6)-Methyladenosine co-transcriptionally directs the demethylation of histone H3K9me2. Nat Genet 52, 870–877 (2020).

18. Akhtar, J. et al. m(6)A RNA methylation regulates promoter-proximal pausing of RNA polymerase II. Mol Cell 81, 3356–3367 e3356 (2021).

19. Lee, J.H. et al. Enhancer RNA m6A methylation facilitates transcriptional condensate formation and gene activation. Mol Cell 81, 3368–3385 e3369 (2021).

20. Xu, W. et al. Dynamic control of chromatin-associated m(6)A methylation regulates nascent RNA synthesis. Mol Cell 82, 1156–1168 e1157 (2022).

21. Liang, Y. et al. METTL3-Mediated m(6)A Methylation Regulates Muscle Stem Cells and Muscle Regeneration by Notch Signaling Pathway. Stem Cells Int 2021, 9955691 (2021).

22. Kudou, K. et al. The requirement of Mettl3-promoted MyoD mRNA maintenance in proliferative myoblasts for skeletal muscle differentiation. Open Biol 7 (2017).

23. Diao, L.T. et al. METTL3 regulates skeletal muscle specific miRNAs at both transcriptional and post-transcriptional levels. Biochem Biophys Res Commun 552, 52–58 (2021).

24. He, L. et al. CRISPR/Cas9/AAV9-mediated in vivo editing identifies MYC regulation of 3D genome in skeletal muscle stem cell. Stem Cell Reports (2021).

25. Machado, L. et al. In Situ Fixation Redefines Quiescence and Early Activation of Skeletal Muscle Stem Cells. Cell Rep 21, 1982–1993 (2017).

26. Xiang, Y. et al. RNA m(6)A methylation regulates the ultraviolet-induced DNA damage response. Nature 543, 573–576 (2017).

27. Lepper, C., Conway, S.J. & Fan, C.M. Adult satellite cells and embryonic muscle progenitors have distinct genetic requirements. Nature 460, 627–631 (2009).

28. Yesbolatova, A. et al. The auxin-inducible degron 2 technology provides sharp degradation control in yeast, mammalian cells, and mice. Nat Commun 11, 5701 (2020).

29. Su, R. et al. Global profiling of RNA-binding protein target sites by LACE-seq. Nat Cell Biol 23, 664–675 (2021).

30. Patil, D.P. et al. m(6)A RNA methylation promotes XIST-mediated transcriptional repression. Nature 537, 369–373 (2016).

31. Gheller, B.J. et al. A defined N6-methyladenosine (m(6)A) profile conferred by METTL3 regulates muscle stem cell/myoblast state transitions. Cell Death Discov 6, 95 (2020).

32. Chen, Y., Chen, P.L., Chen, C.F., Jiang, X. & Riley, D.J. Never-in-mitosis related kinase 1 functions in DNA damage response and checkpoint control. Cell Cycle 7, 3194–3201 (2008).

33. Elliott, D.J., Dalgliesh, C., Hysenaj, G. & Ehrmann, I. RBMX family proteins connect the fields of nuclear RNA processing, disease and sex chromosome biology. Int J Biochem Cell Biol 108, 1–6 (2019).

34. Heinrich, B. et al. Heterogeneous nuclear ribonucleoprotein G regulates splice site selection by binding to CC(A/C)-rich regions in pre-mRNA. J Biol Chem 284, 14303–14315 (2009).

35. Zhou, K.I. et al. Regulation of Co-transcriptional Pre-mRNA Splicing by m(6)A through the Low-Complexity Protein hnRNPG. Mol Cell 76, 70–81 e79 (2019).

36. Lesbirel, S. & Wilson, S.A. The m(6)Amethylase complex and mRNA export. Biochim Biophys Acta Gene Regul Mech 1862, 319–328 (2019).

37. Chenette, D.M. et al. Targeted mRNA Decay by RNA Binding Protein AUF1 Regulates Adult Muscle Stem Cell Fate, Promoting Skeletal Muscle Integrity. Cell Rep 16, 1379–1390 (2016).

38. Hausburg, M.A. et al. Post-transcriptional regulation of satellite cell quiescence by TTP-mediated mRNA decay. Elife 4, e03390 (2015).

39. Figueroa, A. et al. Role of HuR in skeletal myogenesis through coordinate regulation of muscle differentiation genes. Mol Cell Biol 23, 4991–5004 (2003).

40. Yue, L., Wan, R., Luan, S., Zeng, W. & Cheung, T.H. Dek Modulates Global Intron Retention during Muscle Stem Cells Quiescence Exit. Dev Cell (2020).

41. Yesbolatova, A., Natsume, T., Hayashi, K.I. & Kanemaki, M.T. Generation of conditional auxin-inducible degron (AID) cells and tight control of degron-fused proteins using the degradation inhibitor auxinole. Methods 164-165, 73-80 (2019).

42. Huang, Y. et al. Large scale RNA-binding proteins/LncRNAs interaction analysis to uncover lncRNA nuclear localization mechanisms. Brief Bioinform 22 (2021).

43. Chen, F. et al. YY1 regulates skeletal muscle regeneration through controlling metabolic reprogramming of satellite cells. EMBO J 38 (2019).

44. Zhao, Y. et al. MyoD induced enhancer RNA interacts with hnRNPL to activate target gene transcription during myogenic differentiation. Nat Commun 10, 5787 (2019).

45. Li, Y. et al. Long noncoding RNA SAM promotes myoblast proliferation through stabilizing Sugt1 and facilitating kinetochore assembly. Nat Commun 11, 2725 (2020).

46. Xu, W. et al. METTL3 regulates heterochromatin in mouse embryonic stem cells. Nature 591, 317–321 (2021).

47. Shen, S. et al. rMATS: robust and flexible detection of differential alternative splicing from replicate RNA-Seq data. Proc Natl Acad Sci U S A 111, E5593–5601 (2014).

