## supplemental figures for "Nuclear m6A Reader YTHDC1 Promotes Muscle Stem Cell Activation/Proliferation by Regulating mRNA Splicing and Nuclear Export"

### Suppl. Fig. S1. Qiao Y and Sun Q et. al.

**A** **RNA-seq(FPKM)**

| | | QSC | FISC | ASC-24h | ASC-48h | ASC-72h |
|---|---|---|---|---|---|---|
| Writer | Mettl3 | 13.71 | 7.42 | 18.82 | 18.12 | 16.67 |
| | Mettl14 | 5.31 | 2.09 | 12.21 | 15.57 | 12.43 |
| | Ctrlap | 22.63 | 15.53 | 30.65 | 35.28 | 29.69 |
| | Rbm15 | 1.59 | 8.12 | 5.99 | 6.91 | 4.49 |
| | Rbm15b | 7.84 | 30.06 | 33.07 | 31.45 | 20.14 |
| | Zc3h13 | 21.48 | 5.19 | 16.62 | 16.42 | 15.70 |
| | HAKAI | 4.22 | 4.33 | 7.04 | 7.31 | 6.01 |
| Reader | hnRNPG | 41.91 | 15.86 | 35.05 | 40.87 | 31.44 |
| | Ythdc1 | 39.09 | 82.55 | 27.73 | 33.69 | 32.69 |
| | Ythdc2 | 4.46 | 2.70 | 3.07 | 3.45 | 3.60 |
| | Ythdf1 | 17.57 | 36.19 | 40.04 | 42.53 | 31.64 |
| | Ythdf2 | 9.21 | 14.48 | 21.74 | 26.35 | 17.99 |
| | Ythdf3 | 16.57 | 15.46 | 19.85 | 25.68 | 31.70 |
| | Hnrnpa2b1 | 194.44 | 310.07 | 410.18 | 411.56 | 360.10 |
| | Hnrnpc | 48.87 | 118.41 | 155.42 | 176.90 | 151.02 |
| | Igf2bp1 | 0.01 | 0.02 | 0.15 | 0.04 | 0.10 |
| | Igf2bp2 | 0.15 | 25.83 | 3.11 | 2.31 | 5.22 |
| | Igf2bp3 | 0.34 | 0.29 | 0.26 | 0.38 | 0.44 |
| | Fmr1 | 11.99 | 14.69 | 18.48 | 28.50 | 23.92 |
| Eraser | Fto | 22.99 | 17.92 | 23.25 | 21.11 | 28.39 |
| | Alkbh5 | 12.62 | 15.25 | 25.34 | 24.93 | 26.66 |

**Suppl. Fig. S2. Qiao Y and Sun Q et. al.**

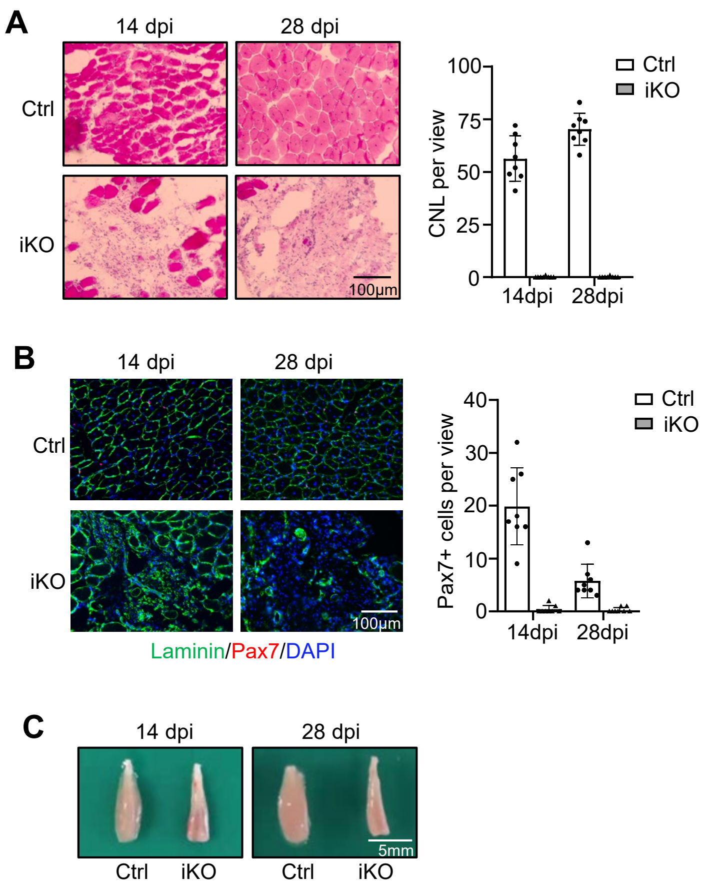

**Suppl. Fig. S3. Qiao Y and Sun Q et. al.**

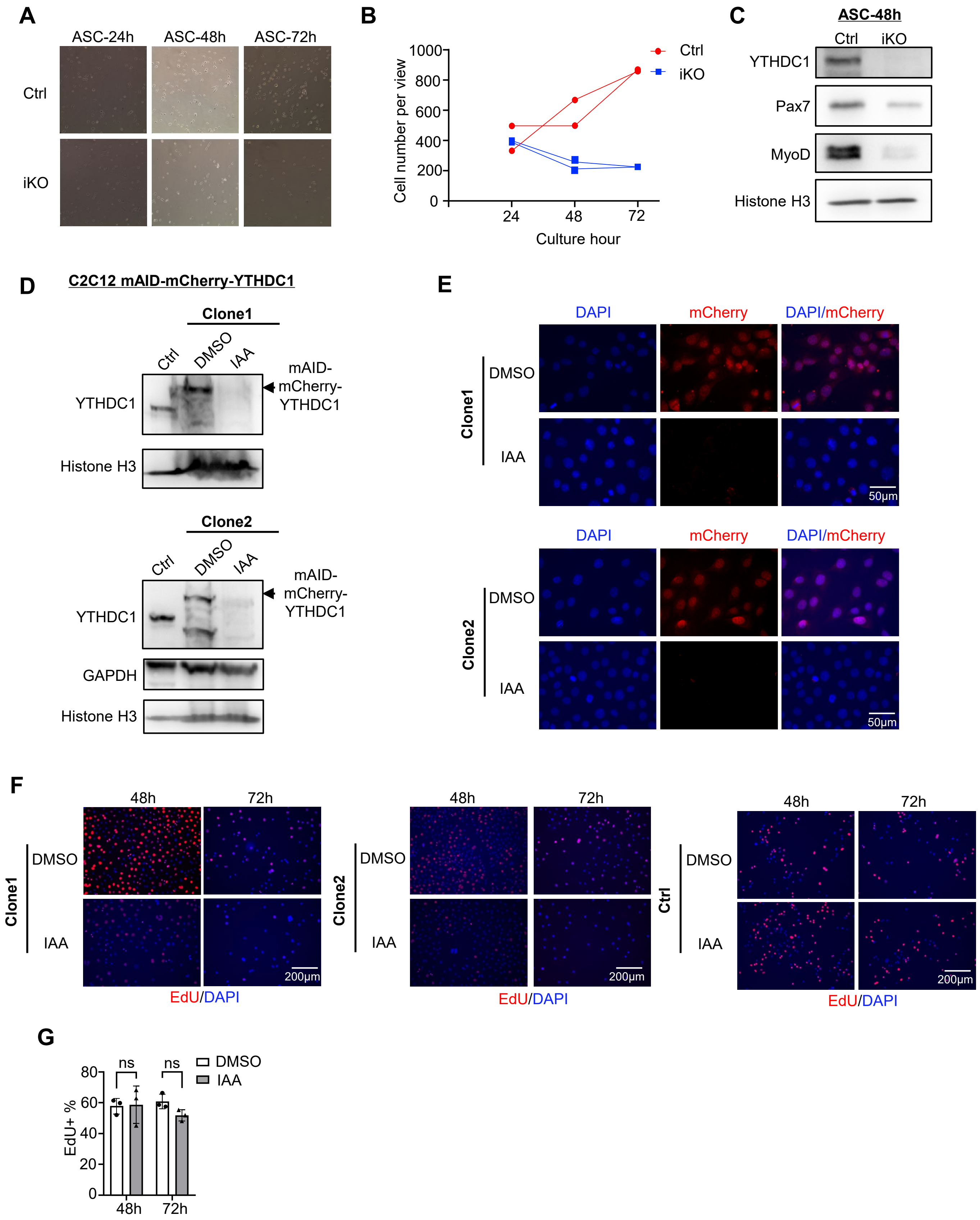

**Suppl. Fig. S4. Qiao Y and Sun Q et. al.**

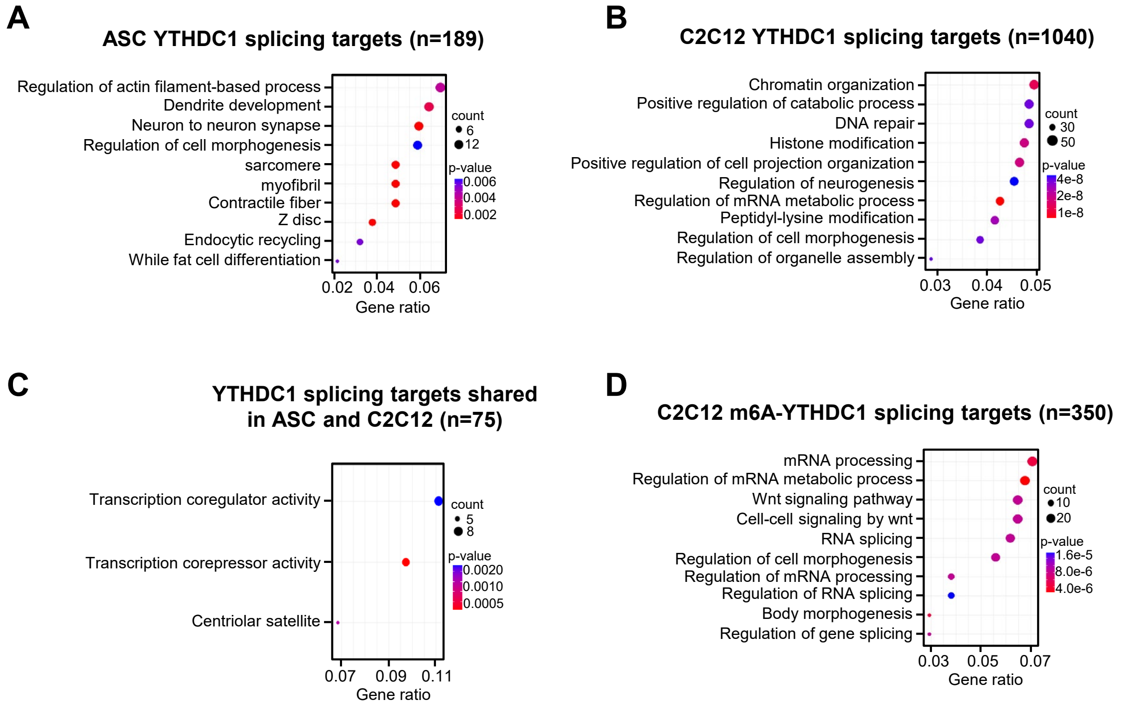

**Suppl. Fig. S5. Qiao Y and Sun Q et. al.**

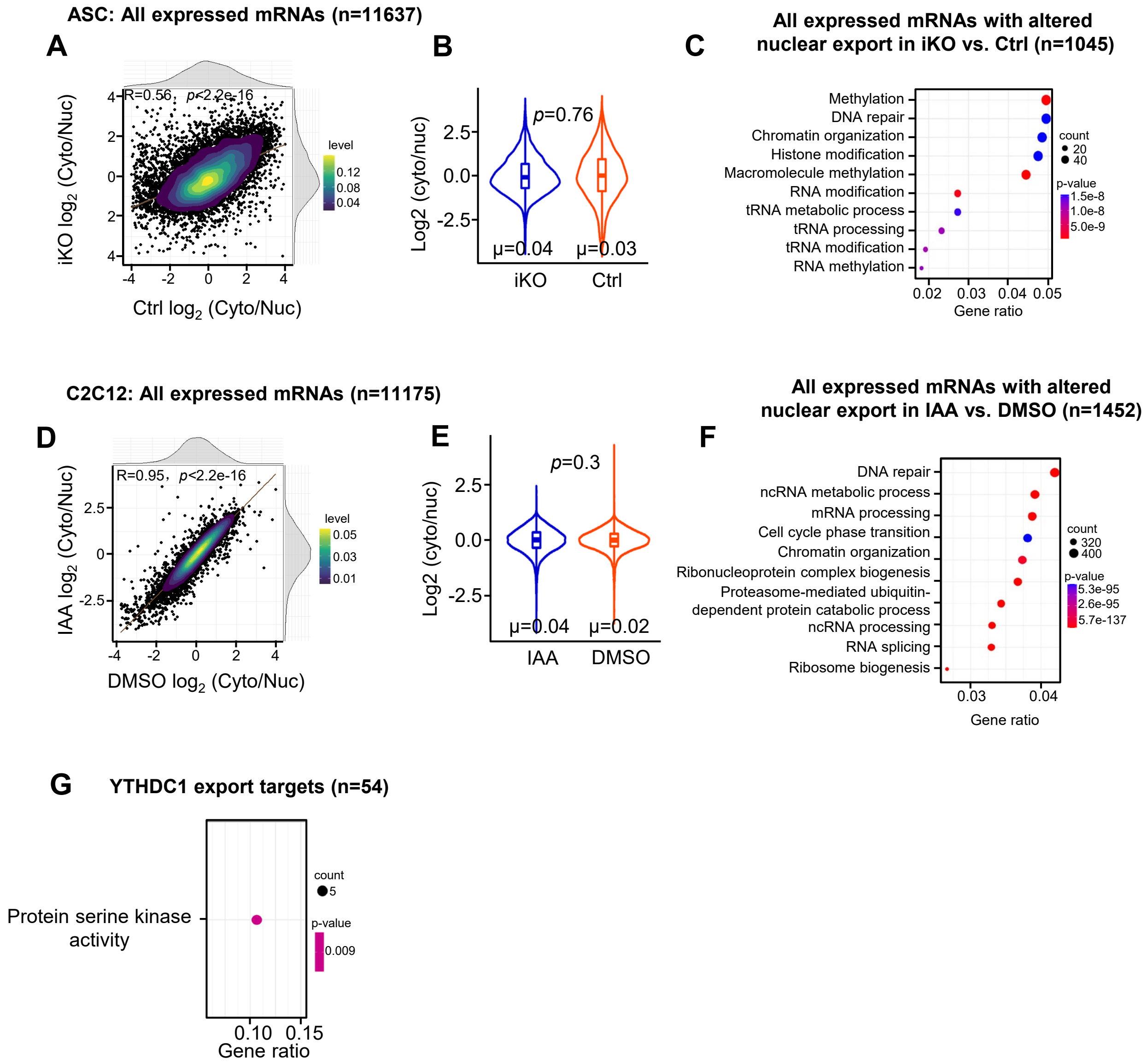
