## supplemental infomation for "Nuclear m6A Reader YTHDC1 Promotes Muscle Stem Cell Activation/Proliferation by Regulating mRNA Splicing and Nuclear Export"

**Inventory of Supplementary Information**

**1. Supplementary Figures**

Figure S1. m6A regulators are dynamically expressed during SC lineage progression and YTHDC1 is induced upon SC activation/proliferation.

Figure S2. Inducible YTHDC1 deletion in SCs abolishes acute injury induced muscle regeneration.

Figure S3. Inducible YTHDC1 knockout impairs SC activation/proliferation.

Figure S4. YTHDC1 depletion in ASCs leads to altered splicing events.

Figure S5. YTHDC1 loss inhibits mRNA nuclear export.

**2. Supplementary Tables**

Table S1. Transcriptomic changes upon YTHDC1 iKO in ASC.

Table S2. LACE-seq profiling of YTHDC1 binding in ASC and C2C12 myoblasts.

Table S3. Analysis of YTHDC1 splicing regulation in ASC and C2C12.

Table S4. Analysis of YTHDC1 mRNA nuclear export in ASC and C2C12 myoblasts.

Table S5. Co-IP/MS identifies YTHDC1 interacting proteins in C2C12 myoblasts.

Table S6. Sequences of oligos used in the study.

**3. Supplementary Figure Legends**

**Supplementary Figure S1. m6A regulators are dynamically expressed during SC lineage progression and YTHDC1 is induced upon SC activation/proliferation.** A. RNA-seq measured expression (FPKM) of m6A regulators during SC lineage progression.

**Supplementary Figure S2. Inducible YTHDC1 deletion in SCs abolishes acute injury induced muscle regeneration. A.** Left: H&E staining of TA muscles from Ctrl and

YTHDC1-iKO at 14 or 28-day post injection (dpi). Scale bar=100μm. Right: Quantification of Centrally Localized Nuclei per view. Bars represent mean ± s.d. of data from 8 views. **B.** Left: IF staining of Pax7(red) and laminin (green) of the above muscles. Scale bar=100μm. Right: Quantification of Pax7+ SCs per view. Bars represent mean ± s.d. of data from 8 views. **C.** Images of the above TA muscles.

**Supplementary Figure S3. Inducible YTHDC1 knockout impairs SC activation/proliferation. A.** ASCs from Ctrl or iKO were cultured for 24, 48 and 72h and light microscopy images are shown. **B.** Quantification of cells per view at the above time points. *n*= 2 mice per group. **C.** Western blotting showing the expression of YTHDC1, Pax7 and MyoD in ASC-48h from Ctrl and iKO. Histone H3 was used as a loading control. **D.** Western blotting showing the degradation of YTHDC1 after treating mAID-YTHDC1 clone 1 or 2 with 1μM 5-Ph-IAA (IAA) for 8hr. DMSO treatment was used as a control. **E.** The above YTHDC1 degradation was confirmed by examining mCherry signal after treating the cell with 1μM IAA or DMSO for 24h. **F.** EdU staining of clone1, clone2 and control cells treated with DMSO or IAA for the designated time points to show the reduced cell proliferation upon YTHDC1 degradation. C2C12 expressing Ostir1 (no AID knock-in) was used as control (Ctrl) to show that IAA small molecule does not affect cell proliferation. **G.** Quantification of Edu+ percentage of C2C12 (without AID knock-in) treated with DMSO or IAA for designated time. Bars represent mean ± s.d. for three views.

**Supplementary Figure S4. YTHDC1 depletion in ASCs leads to altered splicing events. A.** GO analysis of YTHDC1 splicing targets in ASC**. B.** GO analysis of YTHDC1 splicing targets in C2C12 myoblasts. **C.** GO analysis of shared YTHDC1 splicing targets in ASC and C2C12. **D.** GO analysis of m6a-YTHDC1 splicing targets in C2C12.

**Supplementary Figure S5. YTHDC1 loss inhibits mRNA nuclear export. A.** Subcellular RNA-seq was performed using cytosolic and nuclear fractions isolated from ASC-48h of Ctrl and YTHDC1 iKO. The log2 (cyto/nuc) expression change was calculated for all expressed mRNAs. On the top and right, the density plot of log2 (cyto/nuc) expression changes is depicted. **B.** Quantification of log2(cyto/nuc) changes for the above mRNAs in iKO vs. Ctrl. in iKO vs Ctrl. **C.** GO analysis for the above mRNAs. **D.** The above assay/analysis was performed in DMSO or IAA treated mAID-YTHDC1 C2C12 myoblasts. The log2 (cyto/nuc) expression change was calculated for all expressed mRNAs. On the top and right, the density plot of log2 (cyto/nuc) expression changes is depicted**. E.** Quantification of log2(cyto/nuc) change of the above mRNAs upon YTHDC1 degradation. **F.** GO analysis for the above mRNAs**. G.** GO analysis of YTHDC1 export targets in ASC**.**
